# A flexible micro-piston device for mechanical cell stimulation and compression in microfluidic settings

**DOI:** 10.1101/2021.01.11.426203

**Authors:** Sevgi Onal, Maan M. Alkaisi, Volker Nock

**Affiliations:** Electrical and Computer Engineering, University of Canterbury, Christchurch, New Zealand MacDiarmid Institute for Advanced Materials and Nanotechnology, Wellington, New Zealand; Biomolecular Interaction Centre, University of Canterbury, Christchurch, New Zealand

**Keywords:** Microfluidics, Mechanical actuation, Polydimethylsiloxane micropiston, Cell compression, Cancer biomechanics, SKOV-3

## Abstract

Evidence continues to emerge that cancer is not only a disease of genetic mutations, but also of altered mechanobiological profiles of the cells and microenvironment. This mutation-independent element might be a key factor in promoting development and spread of cancer. Biomechanical forces regulate tumor microenvironment by solid stress, matrix mechanics, interstitial pressure and flow. Compressive stress by tumor growth and stromal tissue alters the cell deformation, and recapitulates the biophysical properties of cells to grow, differentiate, spread or invade. Such a solid stress can be introduced externally to change the cell response and to mechanically induce cell lysis by dynamic compression. In this work we report a microfluidic cell-culture platform with an integrated, actively-modulated actuator for the application of compressive forces on cancer cells. Our platform is composed of a control microchannel in a top layer for introducing external force and a polydimethylsiloxane (PDMS) membrane with monolithically integrated actuators. The integrated actuator, herein called micro-piston, was used to apply compression on SKOV-3 ovarian cancer cells in a dynamic and controlled manner by modulating applied gas pressure, localization, shape and size of the micro-piston. We report fabrication of the platform, characterization of the mechanical actuator experimentally and computationally, as well as cell loading and culture in the device. We further show use of the actuator to perform both, repeated dynamic cell compression at physiological pressure levels, and end-point mechanical cell lysis, demonstrating suitability for mechanical stimulation to study the role of compressive forces in cancer microenvironments.

## 1 Introduction

Extensive efforts have been made to study the role of gene mutations in cancer and an accumulation of multiple mutations has been proposed as being necessary for cancer development. Recently however, evidence has been accumulated indicating that cancer is not only a disease of genetic mutations, but that the micro- and nano-environments of cells may be essential factors in triggering tumor growth [1]. For example, the dysfunctional collagen crosslinking in the extracellular matrix (ECM) have been found to lead to breast tumorigenesis and modulate the ECM stiffness to force focal adhesions, integrin expression and, in turn, breast malignancy [2]. Inevitably, tumors can initiate due to an induction either from tumor microenvironment or genetic and epigenetic background of the cells [1]. However, metastasis is the main cause of deaths in cancer patients, thus recent research is elaborating on the basis of metastasis and potential treatments [3]. Tumorigenic and metastatic events are induced by mechanical forces from the altered cell and ECM mechanics [1, 4–6]. Biomechanical forces regulate tumor microenvironment by solid stress, matrix mechanics, interstitial pressure and flow [4]. Cancer cells alter their own biophysical properties and exert physical forces during primary tumor growth and then to spread, invade or metastasize [6, 7]. All these examples point to that phenotype may become dominant over genotype of the tumor cells depending on the microenvironment [1]. Thus, an altered mechanobiological profile of the cells and microenvironment (a mutation-independent element) is proposed to be necessary to promote development and spread of cancer [8]. However, there is no coherent quantitative data on the nature and level of mechanical forces that influence the interactions between the physical micro- and nano-environment and cancer cells [7].

As a result, the application of mechanical compression on living cells, such as cancerous [5, 9–19], non-cancerous and stromal cells [20–22], neurons [23] and chondrocytes [24], has gained importance in recent years. Compression applied on cancer types, such as breast [5, 16, 18, 19], brain [13], pancreatic [9] and ovarian [17, 25] cancer cells, resulted in more invasive and metastatic forms. While indicative, previous studies all point out the need for further investigations to understand the effect of compressive mechanical stimuli in metastasis of different cancer types, for example ovarian cancer [17, 25, 26]. Ovarian cancer cells are exposed to compressive stress mainly by tumor growth, native tissue and hydrostatic pressure from the ascites [25, 26]. SKOV-3 is among the ovarian cancer cell lines tested to be responsive to compressive stimuli and has metastatic phenotype. Thus, SKOV-3 cells can be used as a model to understand the impact of the compression in epithelial ovarian cancer with intraperitoneal metastatic dissemination induced by direct extension of cells and multicellular aggregates into the peritoneal cavity [27]. Understanding these compressive forces may eventually help with the development of pharmaceuticals for the signal transduction mechanisms associated with the mechanical stimulation, [9] and mechanical treatment [12, 28] or mechanoceuticals.

From a cell-biology perspective, mechanical compressive stress can induce the necrosis mode of cancer cell death, resulting in autolysis. Such phenomena play important roles in the recruitment of immune cells to the site of cancer through the release of danger-associated molecular patterns during necrosis [12]. Compression also alters the cell deformation and causes mechanical lysis of the cells. In general, controlled mechanical lysis of adherent cells *in vitro* has advantages over chemical lysis by lytic agents or electrical lysis by applied electrical fields. Intracellular contents can be retained in the sample after mechanical lysis and rapid cell-based assays can be run after compression [10]. Compressive force induction *in vitro* further allows to study stromal cell differentiation and their activation to cancer-associated form (e.g. fibroblast--to-myofibroblast differentiation) depending on the magnitude of the applied stress [4, 22]. Controlled compression in particular can be readily achieved in microfluidic settings. Microfluidic systems can be designed to have integrated physical structures introducing static and dynamic physical inputs, as well as gradients, and enabling real-time imaging, thus providing very useful tools to study biomechanics of the living cells.

While force application systems capable of inducing compressive stress on living cells exist in literature [5, 10, 23, 24, 29], only a very limited number have yet been used to apply compression on cancer models in a localized, flexible and dynamic manner. Tse *et al*. developed a transmembrane pressure device applying compression on cancer cells; however, this is a bulk system that can apply constant force, lacking automation of modulation of the applied pressure [5]. Similar static setups in bulk systems were used more recently by Kim *et al*. [18] and Li *et al*. [19] on cancer associated stromal cells. Although compression effects on other cell types, such as fibroblasts, neurons, chondrocytes, have been shown using microfluidic devices, compression effects in cancer on-chip need more investigation. Lee *et al*. mentioned dynamic compression; however, their study did not expand on the dynamic compression capability of their devices [24]. While crucial to the latter, retraction and position recovery of the compressed compartment of the device were not shown. Similarly, none of the previous studies demonstrated well-defined pressure supply and sensing methods for automation, readability and portability of the pressure application, or showed live compression on cells in real-time.

Asem *et al*. [25] and Klymenko *et al*. [27] applied static compression at ~3 kPa and at 3.18-3.53 kPa, respectively, on ovarian cancer models in off-chip settings. On the other hand, a recent work by Novak *et al*. remains the only study to date using a bioreactor device, which is yet at millimeter scale, to investigate the effect of compressive mechanical stress on ovarian cancer [17]. OVCAR ovarian cancer cells were exposed to both static and cyclic compression, leading to increases in proliferation, invasive morphology and chemoresistance. When compared to cyclic loading, static compressive stimulation enhanced the aspect ratio of the OVCAR3 cells, but no difference in cellular proliferation could be observed. This results is of particular interest, as chronic mechanical loading has been postulated to aid in ovarian cancer progression, forming a positive feedback loop [26]. Clearly, the effects of cyclic compression require further study and applied cyclic pressures need to be expanded from the range of 3.9 to 6.5 kPa used by Novak *et al*. [17] to the physiologically-relevant 3.7 to 18.9 kPa and above estimated to occur in human tumors [26, 28].

Our system achieves the flexible, localized and dynamic control of such compression by use of a polydimethylsiloxane (PDMS) actuator monolithically fabricated as attached to a membrane. The integrated flexible actuator, herein called micro-piston, can be used to apply compression on cells in a dynamic and controlled manner by modulating amount and duration of the applied gas pressure; shape and size of the actuator; and its localization at x-y-z plane as suspended in the microchannel (Fig. 1). Localized compression of the culture can be achieved by applying compression on cells under the micro-piston as test group, while leaving the cells around the micro-piston non-treated as control group (Fig. 1(a)). This provides the flexibility of having the test and control group in the same microchannel with a spatial and temporal control on the groups. For instance, the same device can be first control and then both test and control by applying dynamic compression, as well as that stains or antibodies can be loaded to do further analysis on the cell groups at the same time. Further, the dimensions of the compartments do not rely on each other by resist master fabrication, as each PDMS layer is fabricated separately and then assembled together (see Fig. S1). Thus, when a compartment of the micro-piston device needs to be changed, it is enough to change dimension of one layer while keeping the others as the same.

**Fig. 1.**
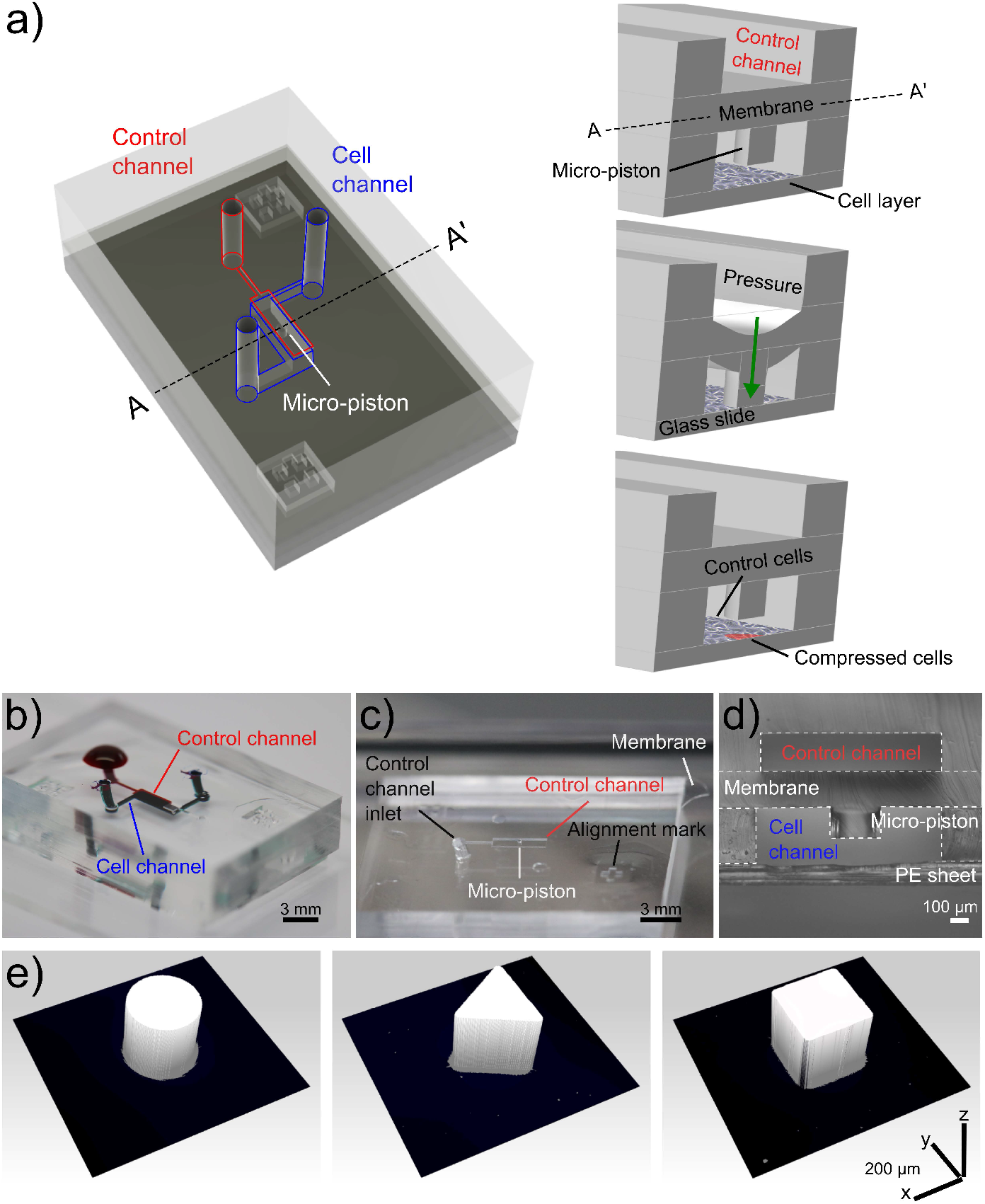
Device design, concept of compression on cells and fabricated devices. (a) Assembly design of the chip composed of control channel (outlined in red) for introducing gas pressure and cell channel (outlined in blue) for cell culture. The concept of applying compression on cells is illustrated by the membrane deflection and micro-piston brought onto the cells by the pressure applied through the control channel and retracted back after compression. Thus, the localized part of the cell monolayer under micro-piston was compressed while the rest acted as control (Figure not drawn to scale). (b) Top view of the PDMS layers of the micro-piston device (Scale 3 mm). The devices were vacuumed for 3 hours and channels were passively filled with dye solutions for visualization. (c) Top view of the PDMS micro-piston with the membrane bonded to control layer before bonding to bottom open channels (Scale 3 mm). (d) Cross-section view of the micro-piston device (Scale 100 μm). (e) 3D profilometer scans of circular, triangle and square micro-pistons sitting on PDMS membranes bonded to control channel layers as in (c) (x-y-z scale 200 μm each).

In this study, we report the development of the microfluidic platform, detailed fabrication method and mechanical characterization of the actuation mechanism by optical imaging methods and computational simulations. We further show evaluation of the platform with SKOV-3 ovarian cancer cells for mechanical stimulation and lysis under dynamic compression. Our results demonstrate suitability of the micro-piston device for mechanical stimulation with various physiological pressures to study cell-biomechanics and compressive forces in cancer microenvironments.

## 2 Results and discussion

The phenotype of cancerous cells changes in response to applied static and dynamic compressive stress. Parameters such as cell growth, morphology and viability are all affected by mechanical stimulation and compression. Compression is unique for ovarian cancer and caused by several primary sources. Both, growth-induced stress from the aberrant cell proliferation, displacing the native cell populations, and external stress, stemming from the native tissue, have been identified as noteworthy contributors to the compressive stress [26, 28]. Moreover, hydrostatic pressure by the excess fluid and ascites expose ovarian cancer cells to additional compressive forces [30], although this factor, depending on the volume of ascitic fluid, largely varies among ovarian cancer patients. Collectively, these stresses evidence that compression has a paramount importance in shaping the mechanobiology of ovarian cancer. To study the role of compression in detail, compressive stress needs to be applied to cells in a localized, flexible and controlled manner. The LOC platform introduced in this work achieves this by use of a PDMS membrane actuator with attached micro-piston. PDMS is well suited for mechanical actuators due to its high elasticity and the very low drift of its properties with time and temperature [31]. Our platform is composed of a control microchannel in a top layer for introducing external force and PDMS membrane with monolithically integrated micro-pistons suspended in a bottom microchannel. This piston was used to apply mechanical compression on ovarian cancer cells cultured on the glass surface enclosing the bottom layer (Fig. 1). Before use with cells, mechanical parameters of the device were characterized via piston actuation to ensure repeatability of compression. This was followed by optimizing the culture of cancer cells in the micro-piston device. Finally, the device was used to study compression at physiologically-relevant pressure levels and mechanical lysis of cells.

### 2.1 Characterization of the micro-piston actuation

Characterization of PDMS device geometries, membrane deflection and thus micro-piston actuation was performed by 3D profilometry and confocal microscopy (Fig. 2). For 3D profilometry, the devices were not covered by a glass surface to enable formation of the optical interference on PDMS surfaces. Also, no cells and media were included in the measurements. For confocal microscopy, the devices were bonded to glass, and bottom channels with micro-pistons were loaded and stained with DiD solution. The confocal measurements were performed either with or without cells and their culture media. Vertical displacement was measured as a function of applied pressure for membranes of different thicknesses. These were obtained by varying PDMS spin-coating and vacuum conditions to better understand the impact process variability may have on piston actuation and ultimately, cell compression. We found that the length the PDMS mixture was kept under vacuum after mixing affected the thickness of the membrane, in particular when slower spin-speeds were used (see Fig. S4). Membrane thickness increased observably for spin-speed of 500 rpm when the vacuum time after mixing was extended from 35 to 90 and 150 min. This effect was not observed for the higher spin speed of 1000 rpm, even after 150 min of degassing, which indicates a correlation with the pot life (the time required for viscosity to double after mixing the base and curing agent) of Sylgard 184 being >120 minutes [32]. It is also important to note that when the vacuum duration after mixing was kept as little as 25-35 minutes, not all of the micro-pistons could be recovered from the 200-μm height resist master on a 4-inch Si wafer due to insufficient degassing time. In contrast, a 100% yield could be regularly obtained when spin-coating at more than 1-hour vacuum durations from the time of mixing.

**Fig. 2.**
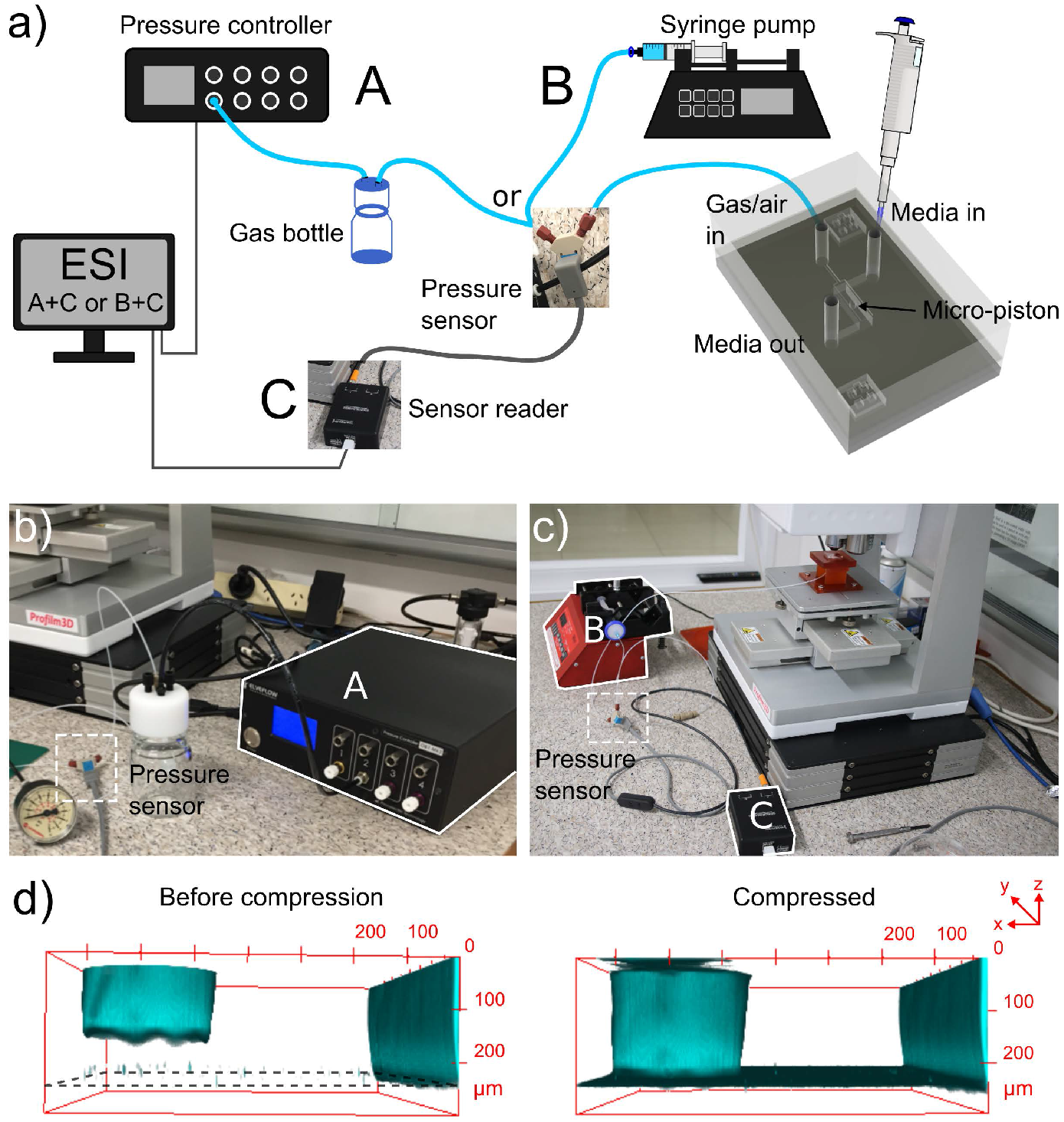
Pressure supply, sensing setup and confocal imaging of the micro-piston device compression (a) Schematic of the experimental setup showing the ways of pressure supply into the control channel and cell seeding. Cells within media were introduced into the cell channel through the inlet labeled as ‘Media in’, and gas or air pressure was supplied through the inlet of the control channel labeled as ‘Gas/air in’. The pressure sensor was located just before the gas flow into the control channel. (b) Experimental setup with pressure controller system. (c) Experimental setup with syringe pumping system. (d) Confocal images illustrating the concept of applying compression with an integrated PDMS micro-piston in a microfluidic channel. PDMS was stained with DiD for visualization.

To characterize membrane deflection and micro-piston actuation, vertical displacement measurements, obtained via optical profilometry, were used to quantify the deflection of 100, 215 and 345 μm thick membranes selected from the spin-coatings with thickness range of 102 ±4.3, 211 ±14.8, 343 ±27.4 (mean ±*σ*), respectively, as monolithically fabricated with 300 μm diameter pistons (Fig. S5). The pressure applications and membrane deflections were applied by using two independent pressure systems, pressure controller system (Fig. 2(a-b) and Fig. S5(a-b)) or syringe pumping system (Fig. 2(a and c) Fig. S5(c-d)). By the two application systems, the vertical displacements of the micro-pistons with thicker, 215 and 345 μm, membranes were linear compared to thinner, 100 μm, membrane with 50-mbar or 0.5-cm^3^ air increments up to 250 mbar applied pressure. In essence, this novel micro-piston device allows using a wide range of membrane thicknesses. However, vertical displacement amount depends on the applied pressure magnitudes and in a final shape of the device it is also restricted by the gap between the micro-piston and bottom surface. Additionally, the pressure magnitudes that can be applied and sensed during the micro-piston actuation still relies on the pressure controller capability and readability of the pressure sensors in use. Thus, in design, the bottom channel total height and the micro-piston height determines the gap which also needs to suit cell culture needs. Membrane thickness to be used in the micro-piston device can be selected based on the designed gap and the range of external pressure application and precise sensing. Taking all into account, for the devices used for cancer cell compression experiments in this study, the average membrane thickness was measured as 211 ±14.8 (mean ±*σ*) μm (n = 164 optical profiles from 7 spin-coatings), where the PDMS was kept under vacuum at 650-700 mmHg for 1.5 hour from mixing (including the 1 hour while PDMS is on the resist master) and spun at 500 rpm for 30 s (Fig. S4). In addition to characterizing the dimensional fidelity of fabricated masters and PDMS replicas (Table T2), compartmental heights of the assembled micro-piston devices were measured for channel total height and micro-piston heights before the device was bonded to glass. To facilitate this, the gap between the micro-piston and bottom surface was calculated for each device by measuring the step height from the leveled surface of the sides of the channel to the surface of the suspended micro-piston in 3D optical profiles (see Fig. S2). Values of the difference between the total height and gap showed close agreement with measured micro-piston height values, illustrating the reliability of the measurements and data analysis for the characterization of the compartments of the micro-piston devices.

Next, characterization of the 215 μm thick membrane deflection with applied higher pressures was followed by micro-piston actuation in an open device to demonstrate position recovery. Figure 3(a) shows the actuation sequence used to measure deflection. Pressure loading profiles were applied using either a pressure controller system (Fig. 3(b)) or syringe pumping system (Fig. 3(c)) and membrane and micro-piston position was found to fully recover to the starting position within measurement limits. This demonstrates that our device can be used for automated dynamic compression and cyclic compression with a pressure controller system. Figure 3(c) in particular, shows that, if no pressurized gas supply is available or a more portable solution is required, semi-automated compression can be performed with a syringe pumping system with an almost identical membrane displacement behavior.

**Fig. 3.**
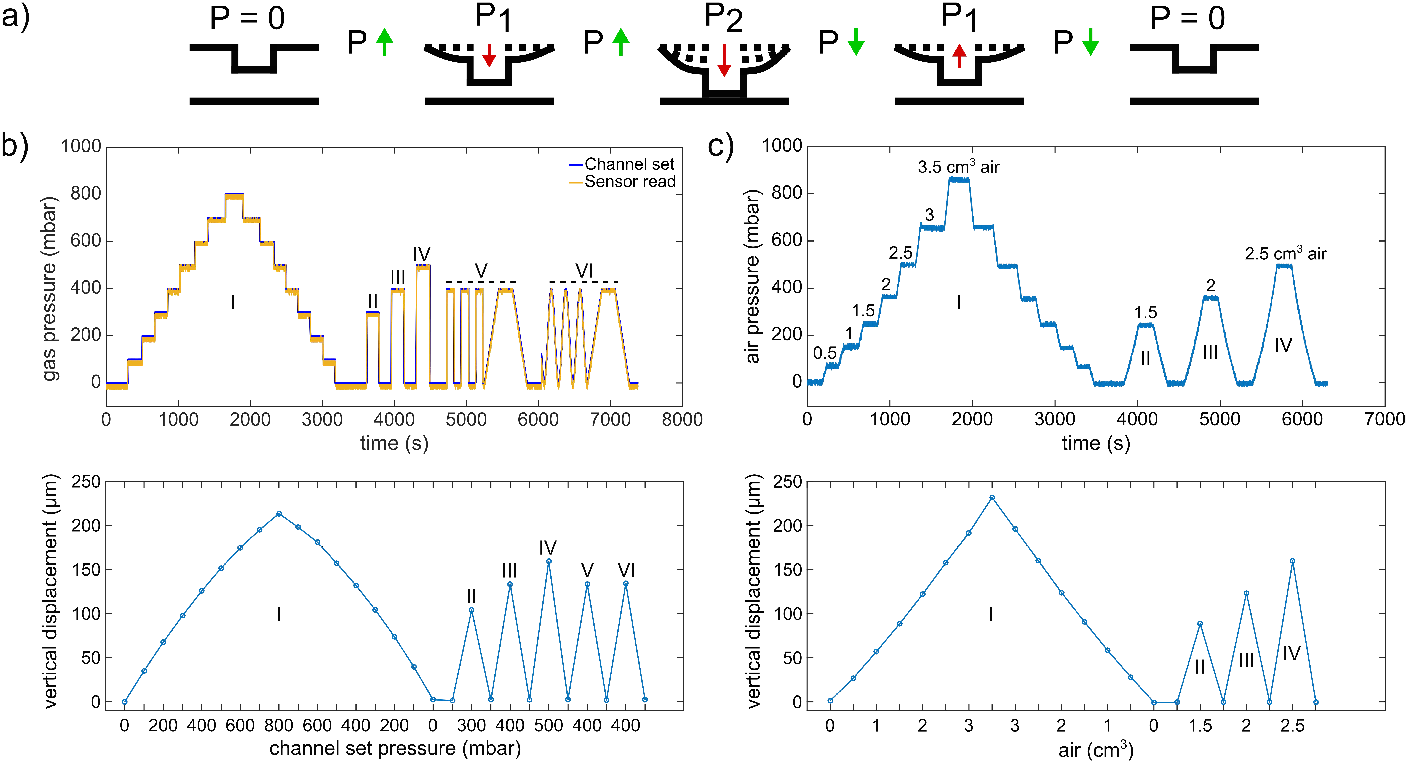
Circular micro-piston actuation using two di?erent pressure application systems. (a) Schematic of membrane deflection and micro-piston actuation by the applied pressure. At static state of the device, there was no external pressure (P), P = 0. Vertical displacement of the membrane and micro-piston was measured while the pressure was kept constant at P = 0, P1 or P2. Membrane and micro-piston were actuated step-by-step by positive (increasing) pressure. Then, the applied pressure was decreased in the same manner of step-by-step and the displacement was measured at each decreased pressure for position recovery. Green arrows indicate when the pressure was increasing or decreasing in between the states. Red arrows indicate the direction membrane and micro-piston moved. (b) Micropiston actuation with various pressure magnitudes and loading profiles (I-VI) for the 215 μm membrane attached to 300 μm diameter piston, generated by pressure controller system. In profile ‘I’, the pressures were positively or negatively applied step-by-step by 100-mbar increments and the displacement was scanned at each step. In profiles ‘II’, ‘III’, and ‘IV’, the pressures were applied suddenly and the displacement was scanned for each sudden pressure amount. In profiles ‘V’ and ‘VI’, cyclic compression where the membrane was deflected and retracted repeatedly was applied by an automated pressure application with (V) or without (VI) time intervals. On the pressure profiles graph, ‘Channel set’ is the pressure that was set through the pressure controller of the system while the ‘Sensor read’ is the pressure that was read through the pressure sensor connected to the gas flow into the control channel of the micro-piston device. (c) Micro-piston actuation with various air amounts leading to corresponding pressure magnitudes in the profiles I-IV for 215 μm membrane attached to 300 μm diameter piston, generated by a syringe pumping system. In profile ‘I’, the pressures were positively or negatively applied step-by-step by 0.5-cm^3^ increments in air amount. The resulting pressures were read through the pressure sensors and the displacement was scanned at each step. In profiles ‘II’, ‘III’, and ‘IV’, the pressures were applied suddenly and the displacement was scanned for each sudden pressure amount.

The concept of applying compression in a fully enclosed device was illustrated using confocal imaging of the PDMS micro-piston stained with DiD cell tracker. This stain was chosen as it allows for simultaneous imaging of PDMS structures and cells [33]. Images were taken before compression and while the micro-piston was compressed with the cell experiment settings, as shown in (Fig. 2d). Staining and confocal imaging were able to visualize the micropiston position inside the device, however, we found the process too slow to capture the dynamic compression on cells, mainly due to the large dimensions of the micro-piston compartment (Fig. S2(a-b)). This result is in agreement with others in literature on confocal microscopy being slow for the real-time volumetric capture of the dynamic compression [24]. Thus, to facilitate the use of wide-field microscopy for imaging the dynamic processes in the device, membrane and micro-piston movement was monitored via the displacement of GFP reporter solution in the piston chamber, as shown in Fig. S3. Again, the same compression profile was applied as in the cell experiment settings and fluorescence intensity change during deflection of the membrane was recorded via time-lapse images. As illustrated by the images in Fig. S3(a), micro-piston touch-down could be tracked (also see Movie V1) and related to device areas where no GFP was present, such as chamber and channel walls. Residual fluorescence signal observed under high pressures (640-1415 mbar, Fig. S3(b)) was attributed to the cells cultured on the glass surface, whereas no cells and thus no GFP was present for the background corresponding to the PDMS sides of the channels. While the vertical displacement of the micro-piston in channel was controlled via the membrane, little change in the GFP signals under the piston was observed for pressures above 640 mbar (Fig. S3(b)). Thus, at higher pressures, the micro-piston was no longer being displaced vertically and the mechanical load was being stored in the compression of the micro-piston itself. On the other hand, the flexible membrane attached to the micro-piston continued to be displaced downwards as evidenced by the GFP solution volume and hence fluorescence signal under the membrane around the micro-piston decreasing with increasing applied pressure (Fig. S3(c)). Compared up to 860 mbar, the pressure values in Fig. S3(b-c), which were read in the fully enclosed devices through the sensory feedback connected within the air flow circuit, were matching the pressure values read during the membrane deflections within the open devices, where the optical profilometry was used (Fig. 3(c)). Optical profilometry and fluorescence solution displacement data was further cross-correlated for three independent devices, as shown in Table T1, demonstrating measurement consistency among devices and between techniques. While recording the GFP images, the focal plane was kept fixed on the cells, as would be the case in cell compression experiments. The average mean gray value ratio obtained in this way was 1.39. If the same experiments were run with images taken at different focal planes of the GFP solution for maximum intensity of a z-stack of fluorescence images, this ratio would be closer to the average height ratio of the devices, which is 1.57. Nonetheless, the data demonstrates that, in lieu of ultra-fast confocal imaging, fluorescent reporter displacement could be used to track piston movement in real-time during cell compression experiments.

### 2.2 Cancer cell loading and culture in micro-piston device

Following mechanical characterization, cancer cell growth and cell viability on the microfluidic platform were investigated by running control and test experiments, either on separate devices or sequentially on a single device. For the optimizations of cell distribution on-chip and cell capture under micro-pistons, devices were loaded with cells either while the micro-pistons were in static state with no external pressure, or driven-up (retracted) towards the control channel with an applied negative pressure of 342 mbar (−342 mbar) as read by the pressure sensor (Fig. S6(a)). Cell capture under micro-pistons was determined to be on average 9 cells per 0.07 mm^2^ for piston-in-static-state loading and increased more than 3-fold to 28 cells per 0.07 mm^2^ for piston-retracted loading. Thus, due to favorable flow conditions in the piston-retracted loading method, more cells were captured under the micro-pistons. Cell distribution on-chip was further analyzed for piston-retracted loading. At zero time of the culture, 11.4% of the loaded cells per field of view (FoV) were under the micro-piston of 14.4% area of the FoV adjusted with the micro-piston at the center (Fig. S6(b) and Table T3). The rest of the cells, 88.6%, were around the micro-piston in the 85.6% area of the FoV. The percentages of cell number under and around micro-piston were not significantly different than the percentages of the corresponding area (p = 0.1417). Thus, homogenous cell distributions could be obtained in piston chamber from the beginning of the cell culture. The cell number under micro-pistons increased on average 83% from Day 0 to Day 1 and 44% from Day 1 to Day 2, providing a confluent cancer cell monolayer under the micro-pistons. Thus, cells grew under and around the micro-pistons at static state of the device regardless the height difference by the gap between the micro-piston and bottom surface compared to rest of the channel (see Movie V2 and Fig. S6(b)).

The lesser gap under the micro-pistons compared to the total height in the rest of the culture channel (Fig. S2) may impact on nutrient supply to the cells. To account for this during mechanical stimulation experiments, cells were kept in their cell-culture growth media to prevent any relative stress that might emerge due to nutrient factors. Moreover, the heights of the cell culture channel (on average 313 μm), and the gap between the bottom of the micropiston and the cells on the glass surface (on average 108 μm) were designed so that cells had access to nutrients in cell-culture growth media at static state by including appropriate cell channel geometries [34]. In particular, the design is expected to allow for passive diffusion of the nutrients over time. In further applications, aim-specific biomolecular gradients can be created from the ports of the bottom layer of the device towards the micro-piston inside the culture channel to study the effect of the generated gradients on cells [35].

As shown in Fig. 4(a), in static culture conditions cell viability was found to be 99.3% in the culture channels, 99.2% in chambers for cells adjacent to the micro-pistons and 99.5% for those under the suspended micro-pistons. These results demonstrate that the device has a cell-safe design and operation until cells under the micro-pistons are compressed by application of high pressures (>300 mbar) to the control layer.

**Fig. 4.**
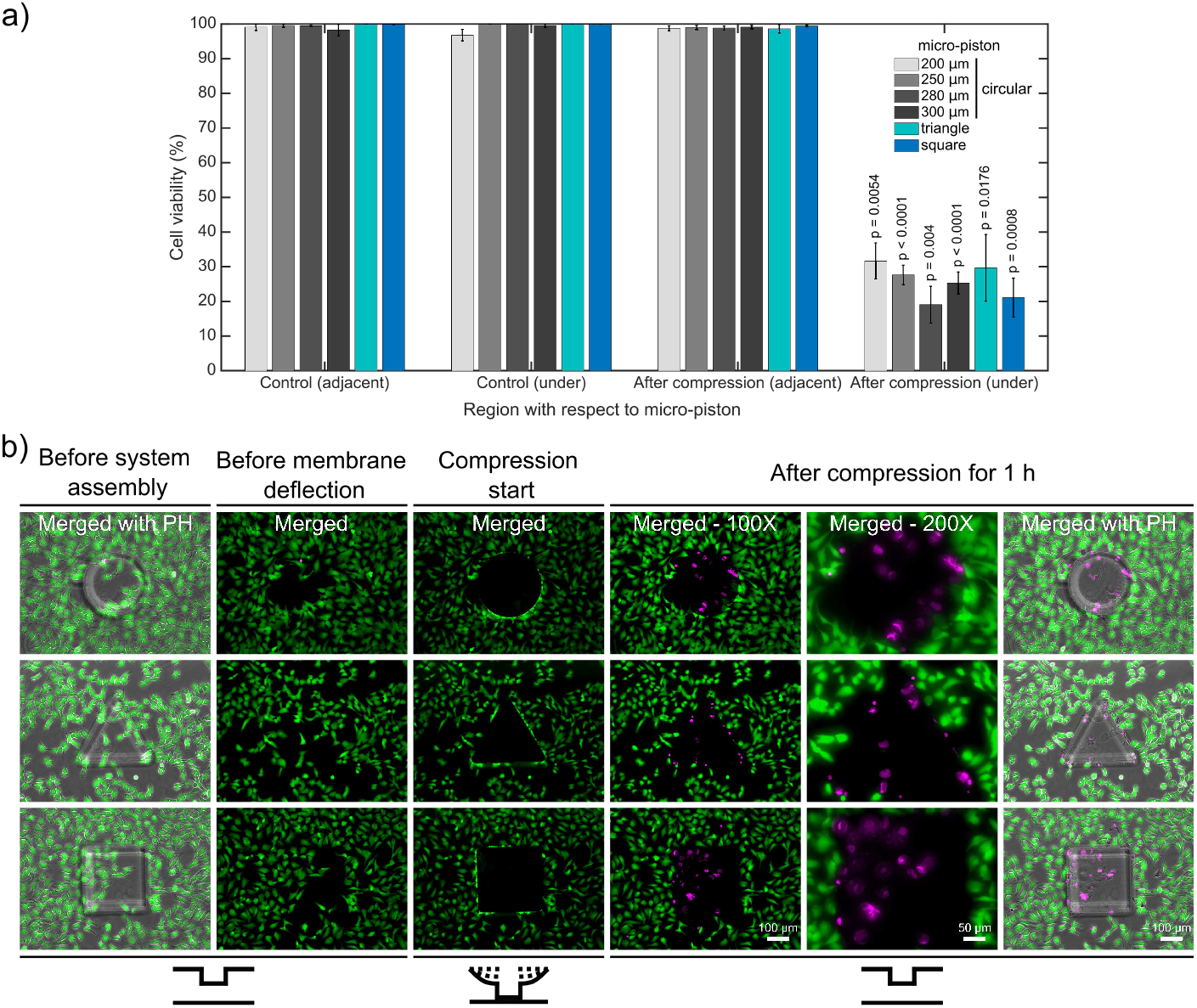
Cell viability for mechanical compression and lysis. (a) Cell viability decreased from on average 99.4% to 25.8% under the micro-pistons of 200, 250, 280 and 300 μm diameter circular, triangle and square shapes (1 h compression, 640 mbar) compared to non-compressed (control) devices (n = 25 devices at static state (non-compressed); n = 25 devices with dynamic compression). The cells within the periphery of the micro-piston, which had part of their body under the micro-piston area, were counted into the group of ‘under the micro-piston’. All other cells in the FoV, outside of the micro-piston periphery, were counted as “adjacent to micro-piston” and accepted as control. (b) Dynamic compression on SKOV-3 cancer cells, mechanical cell lysis and an illustration for patterning the cell monolayer by stamping the particular shape of the micro-piston as circular, triangle or square. Cells and their viability were monitored before the device was connected to the pressure supply unit of the compression system (1^st^column); before membrane deflection (2^nd^column); when membrane was deflected and compression started at 640 mbar (3^rd^column); after compression was applied with different shapes of micro-pistons on SKOV-3 cells for 1 hour and micro-piston position was recovered (4-6^th^columns). After system assembly, and before membrane deflection and compression were started, the cell viability solution was replaced with cell culture media. Live cells retained their Calcein AM signal since the dye entered the cells and was hydrolyzed inside. Once the compression was complete and piston was retracted to its initial position, the cell culture chamber was reloaded with cell viability solution for staining the dead cells. Live cells (green) and dead cells (magenta) were imaged with Calcein AM and EthD-1 epi-fluorescence and merged with phase contrast images.

### 2.3 Compression application and mechanical lysis of the cells

Dynamic application of compression on cell monolayers was demonstrated by deforming and lysing the cells under the micro-pistons and comparing cell viability to control areas on the same device. Cell viability was analyzed for control (non-compressed) devices and after compression with various micro-piston shapes and dimensions. An example of a typical cell compression experiment is shown in Movie V3, where a cell monolayer was first compressed and the micro-piston then retracted. All pressure values during the compression were recorded using a pressure sensor located within the flow circuit before the gas flows into the control microchannel to monitor the micro-piston position feedback in response to the applied pressure. Sensor readings for the micro-pistons were consistent with each other over the length of the experiments and all micro-pistons, including different diameters and different shapes, exhibited reliable position recovery in cell experiments (see Fig. S7).

For in-depth analysis of cell viability on-chip a Calcein AM/EthD-1 cell viability/cytotoxicity kit was used to monitor all the viable and dead cells. Figure 4(b) shows the dynamic compression on SKOV-3 cancer cells and mechanical lysis by the stained cell monolayers in control (non-compressed or before compression) and compressed areas under micro-pistons of different shapes. When the membrane was deflected and the micro-piston moved down onto the cells and burst them by compression, majority of the cells lost their Calcein AM at the beginning of the 1 hour-long compression. Effux of Calcein AM shows that those cells were highly damaged as no fluorescence signal was gained, and hence could not be accepted as viable anymore. The main reason of that the micro-piston was kept compressed for 1 hour is to show how good the device is in operation of the long mechanical compressions, which was also recorded by the pressure sensor in these cell compression experiments (Fig. S7). Cell counting for cell viability analysis on-chip was achieved by optimizing a method of fluorescent label-dependent automated cell counting for adherent cells by dead cell and total cell nuclei staining with Ethidium Homodimer and Hoechst dyes, respectively (Fig. S8). Although cell counting was achieved in parallel with viability testing, by staining both dead and live cells and adjusting the threshold on the images for Calcein AM and EthD-1 signals, respectively, this was further automated by counting the adherent cells via their stained nuclei after validating the correct segmentation of the nuclei.

As shown in Fig. 4(a), in all groups, including control and after compression, cell viability in the channel was calculated for the regions adjacent and under the micro-piston. In doing so, no significant differences in cell viability were observed in control devices between the groups adjacent and under micro-pistons. Additionally, in the regions adjacent to the micro-pistons no significant difference in cell viability in control (non-compressed) and after compression was observed. However, after compression the cell viability under micro-pistons became significantly different (p <0.0001) to that of cells in non-compressed regions in the same device chambers. Cell viability decreased from on average 99.4% to 25.8% under the micro-pistons of 200, 250, 280 and 300 μm diameter circular, triangle, and square shapes after 1-hour compression with 640 mbar gas pressure. At the same time, no significant difference in cell viability was observed for compression between different micro-piston types and sizes. As illustrated by Fig. 4(b), any residual cell viability after compression was mainly due to the cells at the micro-piston periphery, as those cells were being partially compressed, but not necessarily burst by the micro-pistons. Thus, the mechanical compression and lysis of the cells by the micro-pistons in our devices were mainly governed by the applied external pressure, in combination with the thickness of the membrane carrying the micro-pistons. Hsieh *et al*. had varied the diameter of their circular structures (i.e. hydrogel circles under buckled PDMS membrane) from 2 mm to 12 mm and thus changed the compression effect on fibroblast cell alignment [29]. Lee *et al*. had also varied the diameter of their circular structures (i.e. PDMS balloons) from 1.2 mm to 2 mm and observed no significant differences in chondrocyte cell deformation between adjacent diameters (e.g. for 1.2 mm vs. 1.4 mm or 1.4 mm vs. 1.6 mm). They also observed no differences in cell viability between the control (without dynamic compression) and dynamically compressed chondrocytes [24]. Comparatively, in our study, the diameter range has been further downscaled to from 200 μm to 300 μm of circular micro-pistons. If required, changes to the aspect-ratio of the micro-pistons could be used to alter the compression process and hence, cell viability and biological responses of the cells.

Although the cell viability dyes are considered non-toxic at the low concentrations used, [36] we also performed experiments to control for any possible effect of live/dead cell staining and phototoxicity might have besides the mechanical compression. To do so, we compressed the cells after staining them with cell viability dyes and checked the viability during the dynamic compression experiments (see Fig. 4(b)) while, on the other hand, we directly compressed non-stained cells and checked for the viability after the compression (see Fig. S9 and Fig. S10). Results showed that, at 2 μM Calcein AM and 4 μM EthD-1 concentrations, there was no significant effect of the dyes on the cells which were also treated with the mechanical compression (p >0.05). This demonstrates that the cell damage and lysis in our results were mainly caused by the applied mechanical compression. Mechanical lysis is reviewed to be advantageous over the other lysis methods for its high throughput and higher efficiency in lysing cells [37]. In this regard, our platform can apply to controlled release of the intracellular components such as nucleic acids and proteins by adjusting the degree of the mechanical disruption of the cell membranes under the micro-piston while also leaving the control cells around as intact.

Furthermore, we computationally modeled the actuation of the micropiston device, where we established a good agreement of the simulations with the experimental data obtained from the vertical displacement measurements of the micro-piston with pressure controller and syringe pump systems (Fig. 5(a)). This model was going to guide us on predicting the actual pressures inside the device based on the externally applied pressures (Fig. 5(b)). We experimentally conducted dynamic cell compression to establish to which extend cells were affected by the amount of applied pressure and the application time. For this, we applied a sequence of cyclic compression profiles on SKOV-3 cancer cells at mild and then higher pressures in a continuous manner (Fig. 6 and Movie V4). The pressure controller system was used for automation of this cyclic compression application. First, a pressure amount ladder was formed up to 354 mbar, during which the cells showed distinct deformation. Subsequently, cells were set to being chronically compressed for 5 minutes at 354 mbar, followed by a rest for 5 minutes at 0 mbar, as indicated by the steps ‘1b-i, 2b-h’ in Fig. 6(a). This cyclic compression was applied for a total of 1 hour. As can be seen from the image sequence in Fig. 6(a), cells were physically deformed under the micro-piston when being cyclically compressed throughout the cycles ‘1a-i’ (cells at rest) and ‘2a-h’ (cells compressed) (also see Movie V4). The morphology of the compressed cells changed in response to micro-piston contact, with the nuclei and membrane expansion observably different to that of the state before compression at ‘1a’ of Fig. 6(a). In the following compression profiles shown in Fig. 6(b), the application was continued with increasing pressures from 370 and 400, to 640 mbar, respectively. Cells were compressed for up to 2 minutes at each of the higher pressures, which was much shorter than the compression times used in Fig. 4. Cell lysis formed at higher pressures, as expected (see Movie V4), and cell viability was monitored 10-15 minutes after every pressure application profile was complete, allowing for an incubation time for the viability assay. Results shown in ‘1i’ of Fig. 6(a) and ‘1j-l’ in Fig. 6(b) demonstrate that cells were mostly alive after the first cyclic compression profile was complete (‘1i’), despite being deformed distinctly throughout the 1-hour cyclic compression. Simulation of the mechanical actuation (see Fig. 5(b), Fig. S11 and Movie V5) indicated that an actuation with 354 mbar gas pressure corresponded to a contact pressure of 15.6 kPa. While the gap of 109 μm between the piston and glass surface was closing, piston contact pressure started at 1.4 kPa by the applied pressure of 325.9 mbar. In next two of additional 1 mbar, it reached physiological values of 3.6 and 4.5 kPa, and then the piston distinctly deformed cells at 15.6 kPa (Fig. 5(b)). This value fell within the physiological ranges estimated for growth-induced (4.7 - 18.9 kPa) and external (3.7 - 16.0 kPa) compression [28]. The latter has been suggested to be a more noteworthy contributor to the total perceived stress [26] and estimated to exceed 20 kPa based on the experimental data from murine tumors [28]. Following this cycle, more cells were permanently damaged, appearing as dead after step ‘1j’ at 370 mbar (contact pressure of 23.8 kPa from simulation in Fig. 5(b)). Other cells with traces of Calcein AM, but no trace of EthD-1 were considered as damaged but alive, while the cells with traces of the both stains were considered as highly damaged. After step ‘1k’ at 400 mbar (contact pressure of 37.8 kPa from simulation in Fig. 5(b)), only few cells remained alive, with the rest dead. These results were expected, as this and following pressure levels exceeded the upper limits of both estimated growth-induced and external stresses. Finally, apart from a few cells at the micro-piston periphery, all cells directly underneath were dead after step ‘1l’ at 640 mbar (contact pressure of 140 kPa from simulation in Fig. 5(b)). These viability results for short compression times at higher pressures complement results of the 1-hour long compression at 640 mbar in Fig. 4. As observed for this overall long compression experiment, cells in control regions around the micro-piston remained alive at all times during application of the cyclic compression profiles (Fig. 6). Responses of the cells under micro-piston to varying applied pressures in ascending order from Mild (15.6-15.9 kPa) to Intermediate 1 (23.8-26.8 kPa) to Intermediate 2 (37.8-51 kPa) to Severe (127.8-140 kPa) were summarized in Fig. 7. Additionally, the vertical displacement from the position that the piston applied mild pressures on cells until severe pressures bursting the cells, shown in the plateau of the main plot in Fig. 5(b)), is matching the height of the SKOV-3 cells, which is typically extending to around 3 μm [38]. Overall, the ability of our platform to simulate cyclic and varying compression profiles shows that it can be used to further mimic the chronic mechanical stimuli the cells are exposed to in ovarian cancer metastasis [26].

**Fig. 5.**
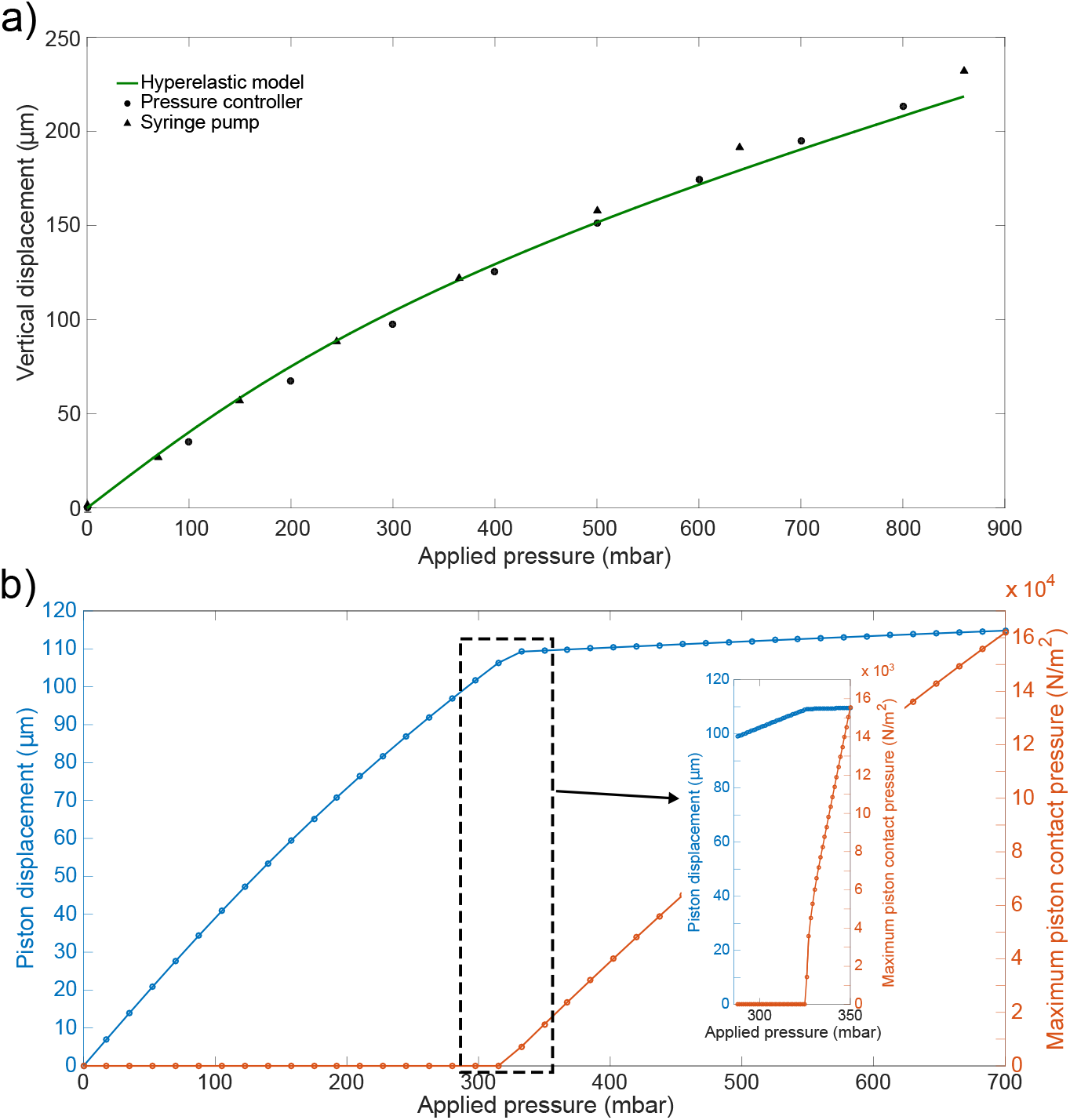
COMSOL simulation results for the actuation of the micro-piston device. (a) Experimental (•, blacktriangle) and computational (line) data comparison. (b) Plot of simulated vertical separation of the micro-piston top and the bottom glass substrate, and maximum contact pressure under the micro-piston as a function of applied gas pressure (boundary load). The applied external pressure was varied from 0 to 700 mbar in 40 steps in the main plot. The inset shows more data points of the initial stage of the piston maximum contact pressure formation at mild pressures, part of the simulation run from 0 to 350 mbar in 320 steps.

**Fig. 6.**
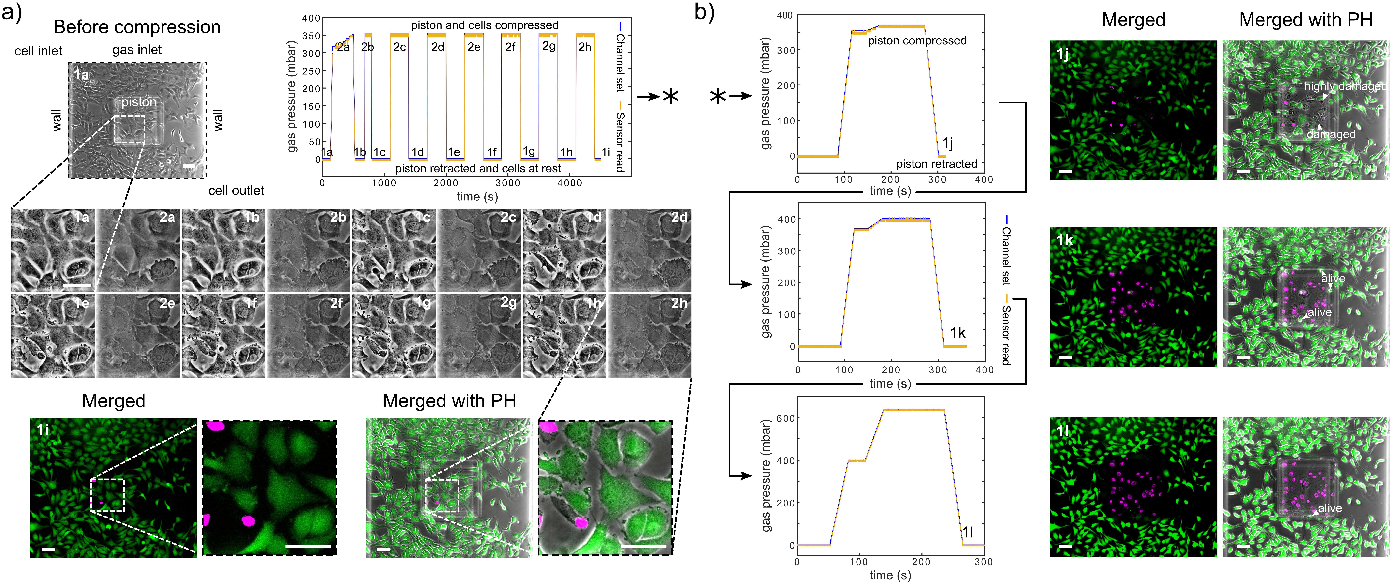
Continuous sequence of cyclic compression profiles applied to a monolayer of SKOV3 cancer cells in a continuous manner. (a) Before compression image of the cells corresponds to step ‘1a’ of the cyclic compression. In the first pressure profile, at step ‘2a’, a pressure amount ladder was formed up to 354 mbar, during which cells showed distinct deformation. Subsequently, cells were chronically compressed for 5 minutes at 354 mbar, then rested for 5 minutes at 0 mbar (1b-i, 2b-h). This cyclic compression was applied for 1 hour. The close-up image sequence shows example cell shapes/deformations under the micro-piston while cells were compressed/rested throughout the cycles, with steps ‘1a-i’ and ‘2a-h’ corresponding to cells at rest and cells compressed, respectively. Cell viability was monitored 10-15 minutes after the pressure application profile was completed (1i). Live cells (green) and dead cells (magenta) were imaged with Calcein AM and Ethidium homodimer-1 epi-fluorescence, respectively, and merged with phase contrast images. (b) Continuation (*) of the pressure application, with pressure increased to 370, 400, and 640 mbar, respectively. Cells were compressed for up to 2 minutes at each higher pressure. Arrow heads (in white) show the representative cells for viability state and cell damage as a result of gradually increasing compression during each corresponding pressure profile. (All scale bars 100 μm).

**Fig. 7.**
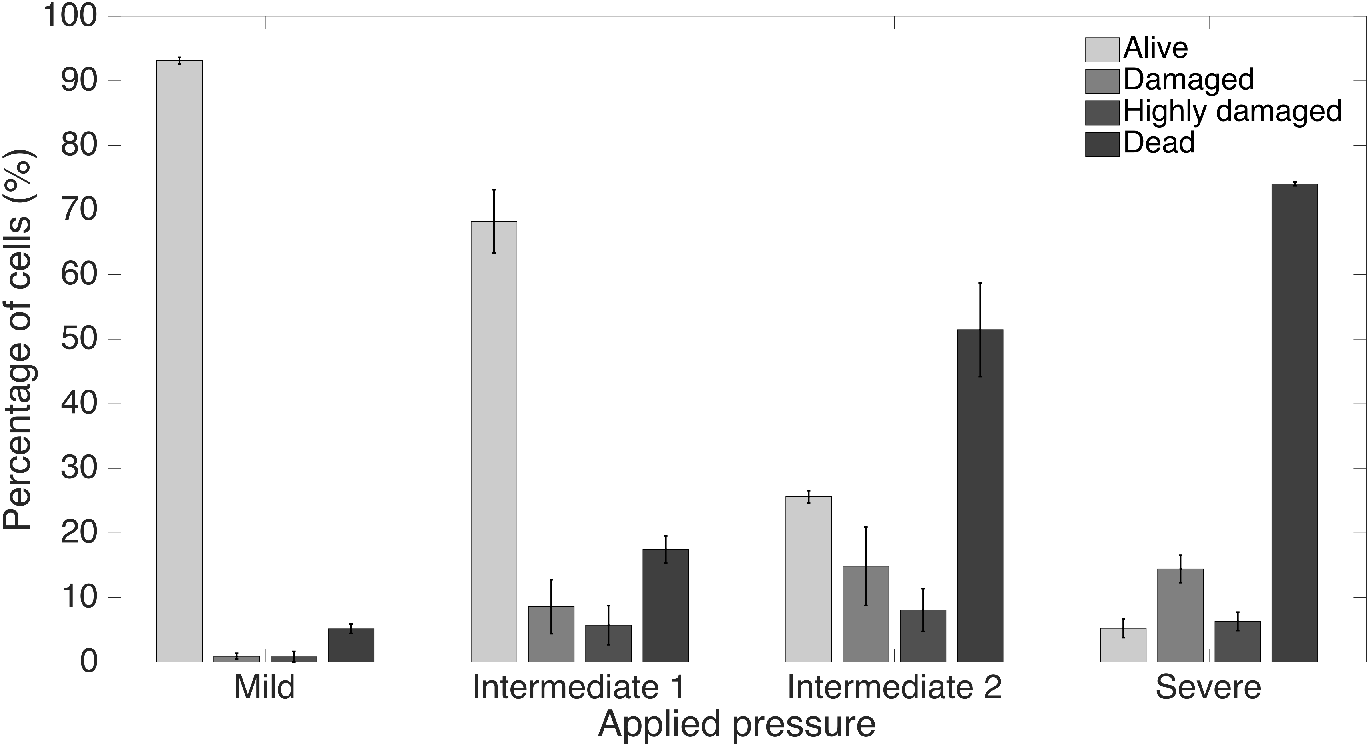
Summary of cancer cell response under micro-piston to varying applied pressures in ascending order from Mild (15.6-15.9 kPa) to Intermediate 1 (23.8-26.8 kPa) to Intermediate 2 (37.8-51 kPa) to Severe (127.8-140 kPa) out of at least 3 independent experiments of cyclic compression using micro-piston devices which were operated at a continuous manner of the xsapplied pressure sequence.

In summary, the platform was able to dynamically compress cell monolayers up to the mechanical lysis of cells at actuation pressures above 300 mbar. By enabling similar experiments to be performed in a more integrated fashion, our study results add to the previous work by Tse *et al*., which showed that mechanical compression can enhance the invasive phenotype of cancer cells and stimulate their migration [5]. While pressure magnitudes obtainable with our system are comparable to those in work by Kim *et al*., who applied up to 35 kPa external compressive stress on MCF7 cancer cells up to lysis [10], the use of micro-pistons for compression in our system adds the advantage of non-compressed regions for direct control on-chip. The importance of control was most recently demonstrated by Lee *et al*., who observed no significant difference in cell viability between compressed and control groups when 14 kPa of pressure was used to mechanically stimulate alginate-chondrocyte constructs [24]. We further showed the capability of the platform to dynamically stimulate cancer cell monolayers with various intermediate actuation pressures in a continuous, cyclic manner with an order of Mild (15.6-15.9 kPa) to Intermediate 1 (23.8-26.8 kPa) to Intermediate 2 (37.8-51 kPa) to Severe (127.8-140 kPa). As such, the results shown here expand on the work by Hosmane *et al*., who used intermediate pressures applied statically by micro-pads to study the mechanics of neuronal cell damage [23]. Their results showed single axon mild injury and regrowth at <55 kPa, moderate levels of injury from 55 to 95 kPa and severe levels of injury for pressures >95 kPa.

Finally, while cell compression with our micro-piston device was shown using ovarian cancer cell monolayer mimicking the impact of compression on direct extension of cells into peritoneal cavity in a metastatic dissemination of epithelial ovarian cancer, the overall device concept can readily be adapted for use with cell-laden hydrogels or by loading a spheroid solution for compression of cells in 3D. The device can be further used in its current form to apply external compression on an endothelial cell monolayer to study the basics of the vessel compression existing in cancer patients, as shown in ex vivo tumor sections [28, 39]. In the future we hope to expand this work to the extended tumor microenvironment including stromal cells, such as fibroblasts and macrophages. These have been reported as sensitive to compressive forces, e.g. fibroblast-to-myofibroblast differentiation, and there are known biochemical interactions between cancer cells and stromal cells, which might be triggered by the solid stress in the microenvironment [4].

## 3 Conclusion

We have introduced a flexible multilayer microdevice with a micro-piston suspended into cell-culture chamber for dynamic mechanical cell stimulation and compression. Fabrication of the device was described and actuation by a monolithic membrane controlled by a gas channel on the top layer was demonstrated. We showed the characterization of the micro-piston actuation using optical imaging methods and two different external pressure system types, adding flexibility based on the laboratory requirements. The actual pressures inside the cell channel were computationally modelled based on the applied pressures and device dimensions. Hence, our device is optimized to apply a whole range of physiological pressure levels, as well as milder and severe pressures. Applicability of the device was demonstrated via the quantification of the response of ovarian cancer cell monolayers to mechanical stimulation in the micro-piston device. Cell viability before and after mechanical compression was used to illustrate the suitability of the platform for cyclic cell compression and lysis. Circular, square and triangular micro-piston outlines and different diameters were successfully propagated into the cell layer.

Based on current studies and future perspectives, our platform provides a useful mechanical tool to study the direct effect of compressive forces on cancer cells and the tumor microenvironment. It can be used to investigate responses of cancerous and healthy cells to applied stress with regards to changes in cell morphology, viability and mechanobiologically-related protein profile. Surface-coating solutions of natural ECM such as fibronectin, laminin, or type 1 collagen can be used to enhance cell adhesion to the culture surface in the microchannels by mimicking the native microenvironments. The platform is potentially suitable for automated creation of wound healing assays on-chip, as the spacing between the micro-piston and cell culture surface, together with the cell number captured under micro-piston can be tightly controlled. Having different micro-piston shapes available will also be of interest in this application and in tissue engineering in general. Overall, our platform constitutes a promising tool for studies of cell-mechanical force interaction, as well as cell-cell, cell-microenvironment and cell-drug interactions, where single or co-cultured cell types need to be patterned and controlled.

## 4 Experimental Section

### 4.1 Device design and fabrication

To demonstrate localized and dynamically controlled compression of cancer cells we fabricated a microfluidic platform with flexible actuators composed of PDMS [40]. Main components of the platform are control channel in the top layer, membrane and micro-piston in the middle, and cell culture chamber in the bottom layer enclosed with glass substrate (see Fig. 1(a)). Control channel was designed to introduce external pressure [41]; membrane to enable micro-piston actuation; and the micro-piston itself to apply compressive forces on the cells cultured on the glass surface in the bottom layer channel. Photolithographic masks were designed using L-Edit (v2019, Mentor Graphics) and written onto photomask blanks (Nanofilm, USA) using a laser mask writer (μ PG101, Heidelberg Instruments). Silicon wafers were dehydrated at 185°C in an oven overnight. After cooling down, the wafers were plasma cleaned at 100 W with O_2_ gas flow at 5 sccm for 10 minutes (Tergeo, PIE Scientific). As shown in Fig. S1(a), dry-film lamination of negative-tone resist on silicon wafers was used to fabricate masters [42]. For the lamination set-up we used 500 μm thick metal shims for a total thickness of Si wafer plus 200 μm or 300 μm SUEX film (DJMicrolaminates) and PET sheet. The films were laminated to the substrate at 65°C at 1 ft/min using a roll laminator (Sky 335R6, DSB). Post-lamination bake was applied at 65C for 15 minutes and the laminates were left to cool down for 2-3 hours before UV exposure on a mask aligner (MA-6, SUSS). Filtered doses (PL-360-LP, Omega Optical) of 1800 mJ/cm^2^ for 200 μm thick film and 2300 mJ/cm^2^ for 300 μm thick film were used in multiple cycles of 20 second exposures with 60 second cool-down intervals between cycles. Post-exposure bake (PEB) was applied at 65°C for 5 minutes and at 95°C for 15 minutes for 200 μm films and 20 minutes for 300 μmfilmswitha ramp up of 100°C/hr and ramp down of 15°C/h. The masters were developed 24 hours after cooling down from PEB. Developer was applied with the SUEX master face-down in Propylene glycol methyl ether acetate (PGMEA, Sigma) until fully developed. Hard-bake was applied at 125°C for 60 minutes with a ramp up of 100°C/h and ramp down of 15°C/h on hotplate.

Before replication, master molds were treated with 1H, 1H, 2H, 2H-Perfluoro decyl triethoxysilane (PFDTES, Sigma-Aldrich) for 3 hours. All flexible layers of the micro-piston device were fabricated from a 10:1 w/w mixture of PDMS (Sylgard 184, Dow) base and curing agent. Micro-pistons, monolithically integrated onto a PDMS membrane, were obtained by spin-coating (Laurell WS-650) PDMS pre-polymer onto 200 μm thick resist masters. After spinning, PDMS was baked for 1 hour at 80°C for initial curing. Similarly, bottom layer channels and the control layer were fabricated by exclusion molding [43], modified for our work, and replica molding of PDMS [44] off 300 and 200 μm thick resist masters, respectively. For the exclusion molding, the PDMS poured on the master was first vacuumed for 2 hours, then a PE sheet was rolled on the PDMS and most of the excess PDMS was excluded out. Finally, a rubber sheet and heavy metal block were placed on the PDMS on master for complete exclusion out of the resist mold and the construct was baked at 80°C for 2 hours, as shown in Fig. S1(a). Left to cool down overnight, the PDMS stamp with open channels and open frame alignment marks was peeled off. Alignment marks designed to fit each other were used to align the layers while bonding (see Fig. 1(a-c) and Fig. S1). After plasma surface treatment at 15 W with O_2_ gas flow at 3 sccm for 1 minute (Tergeo, PIE Scientific), devices were assembled by manual alignment and bonding of the membrane with micro-piston to the control layer and bottom channels (see Fig. 1(c-d)). PDMS layer alignment was checked using a light microscope and, if necessary, corrected for fine matching the edges of top and bottom microchannels before the plasma treated surfaces were permanently bonded. After every alignment and bonding of the layers, the construct was baked for 2 hours at 80°C. These fabrication steps were applied for different diameters and shapes including the circular, triangle and square micro-pistons shown in Fig. 1(e).

### 4.2 Mechanical modelling of micro-piston actuator

Actuation by the micro-piston device as a function of applied gas pressure were modelled as a symmetrical 3D geometry using COMSOL Multiphysics (V5.5, COMSOL). PDMS was modelled as hyperelastic material using a Saint Venant-Kirchhoff model. The Lame parameters λ and *μ* of PDMS were set at 4.66 MPa and 460 kPa, respectively [45]. Borosilicate from the materials library was used to model the glass substrate as linear elastic material. External gas pressure was applied as a boundary load and varied from 0 to 700 mbar in 40 steps using a parametric sweep. The standard stationary solver was used with a physics-controlled mesh. Pressure under the piston was visualized as surface plot, and piston displacement and maximum piston pressure were exported as a function of applied gas pressure.

### 4.3 Pressure supply unit and operation of the force application system

Membrane, and thus micro-piston, were actuated by syringe pump (NE-1000, New Era Pump Systems) or pressure controller (OB1 Mk3+, Elveflow) coupled with pressure sensors (MSP4, 7 bar, Elveflow) and sensor readers (MFR and MSR, Elveflow) operated via ESI software (v3.02, Elveflow Smart Interface) to address the air or N_2_ gas pressure amount required to deflect the membrane and actuate the micro-piston. PDMS devices were fixed through a custom-designed, 3D printed (MoonRay, SprintRay) sample holder stage so that the devices did not move while applying pressure.

### 4.4 Optical profilometer measurements

The membrane deflection and micro-piston movements were quantified using a Profilm3D optical profilometer (Filmetrics Inc., USA) by scanning the total height of the vertical displacement of the deflected membrane and micro-piston during actuation. The data was further processed and analyzed using the ProfilmOnline software. For each applied pressure amount, average step heights of the vertical displacements were calculated from different regions-of-interest (ROIs) on the membrane at each side of the micro-pistons. For the full view of a micro-piston per cell-culture chamber, 3D optical profile series were scanned and stitched on an area of 7 mm height x 9 mm width with 20% overlap between individual scans (see Fig. S2(c)).

### 4.5 Cell culture and preparation

SKOV-3 ovarian cancer cells (supplied by the Laboratory for Cell and Protein Regulation at the University of Otago) were cultured in Earle’s salts and L-Glutamine positive MEM medium (GIBCO^®^) supplemented with 10% fetal bovine serum (FBS, Life Technologies), 1% of penicillin/streptomycin (Life Technologies) and 0.2% fungizone (Life Technologies) in a humidified atmosphere of 5% CO_2_ at 37°C. Cell seeding densities of 0.5M, 1M, 1.5M and 1.8M cells/ml for ovarian cancer cells were tested for optimization of the cell number in microfluidic channel to form a cell monolayer and capture cells under the micro-piston. To promote cell adhesion by enhancing electrostatic interactions between the cell membrane and the glass substrate, the cell culture channels of the microfluidic devices were coated with 0.01% (w/v in H_2_O) poly-L-lysine (PLL) solution (Sigma).

### 4.6 Confocal imaging of the micro-piston device

For confocal imaging of the compression application with the PDMS micropiston, PDMS cell culture channels with the suspended micro-piston inside were stained with DiD (lipophilic carbocyanine DiIC_18_(5) solid) (Vybrant, Thermo Fisher, 1:300) for 48 hours. The DiD solution on-chip was replenished 24 hours after the staining started. The stained micro-pistons and microchannels were imaged using a confocal laser scanning microscope (Leica TCS SP5) with PMT detectors. DiD was excited using the HeNe laser at 633 nm with 46% intensity with an emission at 665 nm. The distance between the bottom glass surface and suspended micro-piston in the cell culture channel was 3D scanned at static state (before compression) and then at compressed state of the micro-piston (see Fig. 2(d)) by taking the outmost upward z-position of the 20X objective (NA 0.70; PL-APO IMM CORR) as reference for the start level of each scan by the physical restraint of the mechanical knob at an inverted microscope setup. The length of each 3D scan was taking up to 5 minutes by 5 μm step-size for a ~300 μm total z-volume with the DiD and brightfield channels at 1024×1024 resolution, due to the dimensions of the micro-piston device. In between the static and compressed states, xyt laser scanning was applied to capture the displacement of the actuated micro-piston while approaching the focal plane of the cells at the bottom surface. Data were processed using the 3D Viewer plug-in of ImageJ (Fiji) [46].

### 4.7 Device operation for cell culture and compression experiments

Before the cell culture on-chip started, micro-piston devices were sterilized under the UV light in a biological cabinet for 30 minutes. Different cell-seeding densities were tested to improve cell density on-chip and the chance of capturing cells under the micro-pistons, as described above. For the homogeneity of cell distribution on-chip and capture under micro-piston, the way of loading cells into the microchannels with those hanging structures was further optimized. Cells were introduced into the culture channels while the micro-piston was either at static state or further driven-up (retracted) towards the control channel. In our piston-retracted method, the micro-pistons were retracted by applying negative pressure by 2 cm^3^air sucked over 4 minutes using syringe pumping system. Cells were introduced into the bottom channel and the micro-piston was kept retracted for 5 minutes to let the cells sink to the PLL-coated glass surface. Then the micro-piston was released to recover its position by applying positive (increasing) pressure. Cells were cultured in the micro-piston devices for at least 2 days and their growth and spread around and under the micro-pistons were monitored and recorded during culture until mechanical compression was applied.

For all samples, compression was applied to cells in culture media to eliminate any non-mechanical stress. The microscopy focal plane was kept on the cells while applying the compression and during movement of the micro-piston towards the cells. Micro-piston movement, compression on cells and mechanical lysis of the cells were recorded with time-lapse imaging. Data of the pressure sensor read was recorded during the compression experiments to monitor the pressure profile in the devices and ensure the devices were working properly. Compression was applied up to 640 mbar over 6 minutes. Cells were kept compressed for 1 hour and the withdraw-rate to 0 mbar and position recovery was the same as the compression rate. For cell viability imaging, the cells were incubated with 2 μM EthD-1 and 4 μM Calcein AM solutions of cell viability/cytotoxicity kit (Invitrogen) for 20 minutes. After the compression experiments, cells were fixed on-chip with 4% Paraformaldehyde (Alfa Aesar) for 30 minutes and nuclei were stained with 1 μg/ml Hoechst solution (Thermo Scientific, 33342) for 20 minutes.

### 4.8 Green fluorescent protein solution displacement by membrane deflection in the micro-piston device

Green fluorescent protein (GFP) was expressed in-house in *E. coli* bacteria. Micro-piston devices were loaded with GFP solution (6.83 mg/ml) and compression was repeated as described for the cell experiments. Micro-piston movement and membrane dynamics were visualized for inference of the physical state from fluorescence intensity change (Fig. S3(a)). Using an inverted microscope, the GFP time-lapse fluorescence images were taken under constant light exposure at 1 minute intervals of the actuation by the applied pressure amounts in Fig. S3(b-c). An A-A’ line was drawn at the region under the micro-piston and the plot profile of the GFP intensity change at each time-lapse frame was shown for the A-A’ line at the applied pressures (Fig. S3(b)). The plot profile for the GFP intensity change under the membrane was shown along a representative line B-B’ on each frame taken at the corresponding pressure amount (Fig. S3(c)) similar to the work of Kim *et al*. on characterization of PDMS membrane deflection with fluorescein [10]. Further, ROIs were drawn at different regions adjacent to the micro-piston under the membrane. Mean gray values were measured from the GFP intensity at those ROIs on the frames taken under the pressures of 0 mbar and 640 mbar for the ratio of the mean gray value at static (non-compressed) to the one at the compressed form (see Supplementary Table T1). This ratio was used for further correlation of the change in GFP intensity with the change in compartmental heights of the micro-piston device during membrane deflections. At least 3 independent experiments with different devices were performed to measure fluorescent intensity change as a result of the displacement of GFP during actuation of the micro-piston in a typical cell experiment setup.

### 4.9 Imaging and data analysis

Images of cells were taken as phase-contrast time-lapse image series using a diascopic and epi-fluorescence inverted microscope (Nikon ECLIPSE Ts2) equipped with a digital camera (Tucsen USB2.0 H Series) and TCapture imaging software. For cell growth, alignment and spread analysis, cells were imaged day-to-day starting from the cell seeding time using 10X (NA 0.25; Ph1 ADL) and 20X (NA 0.45; Plan-Fluor OFN22 Ph1 ADM) objectives. For dynamic cell compression experiments, data recording was started at the same time as the pumping for applying the gas pressure and time-lapse images were captured at every 10 seconds. Cells in the control regions where there was no micro-piston were also imaged before and after compression application. Cell viability images for the cells under micro-piston and at control regions in the channel were taken with EthD-1 and Calcein AM epi-fluorescence.

Recorded images were processed and analyzed using ImageJ (Fiji) [46]. For cell viability analysis, the cells under a micro-piston were determined according to the images with Calcein AM and EthD-1, and cross-checked with the phase-contrast images taken before the deflection and the time-lapse frames recorded during the deflection. With exception of the cells with strong signals of Calcein AM or EthD-1, further analysis was done as follows: If the cell body was under the micro-piston before compression, but totally burst after compression, and had no stain of Calcein AM with or without traces of EthD-1, those cells were counted dead. If any cell had Calcein AM signal and traces of EthD-1 signal at the same time, such cells were counted as damaged but alive, as well as the cells with traces of Calcein AM and without any EthD-1 signal. Video frames recorded during deflection and retraction of the membrane were used to track the cells bursting due to compression, as well as the changes in their morphologies. Cells that were detached from the culture by attaching to the micro-piston surface during compression were counted into the analysis by taking max intensity of the Z-projection of fluorescence images from the glass and PDMS surfaces that came into contact. Two hundred fold magnification images of EthD-1 epi-fluorescence contained the most information for the stained nuclei of the dead cells fitting the region under micro-pistons, and were thus used in the cell viability analysis.

One-way analysis of variance (ANOVA) and Student’s t-test were used to determine significances for the comparisons of cell viability between multiple groups and between two groups, respectively. Two sample t-test between percents (StatPac) was used to compare the percentages of individual groups in on-chip cell distribution data. Statistical significance was taken as p <0.05. Data were represented as mean ±*s.e.m*., except for Fig. S4, in which the data is presented as mean ±*SD*(*σ*) for the PDMS membrane thicknesses measured with the optical profilometer.

## Acknowledgements

We thank Helen Devereux, Gary Turner, and Nicole Lauren-Manuera for technical assistance. We also thank Serena Watkin for gifting the GFP used, James Davies for measurement of GFP concentration and Mathieu Sellier for help with COMSOL. The work was financially supported by the MacDiarmid Institute for Advanced Materials and Nanotechnology and the Biomolecular Interaction Centre.

## Supplementary Information

**Supplementary Figure S1.**
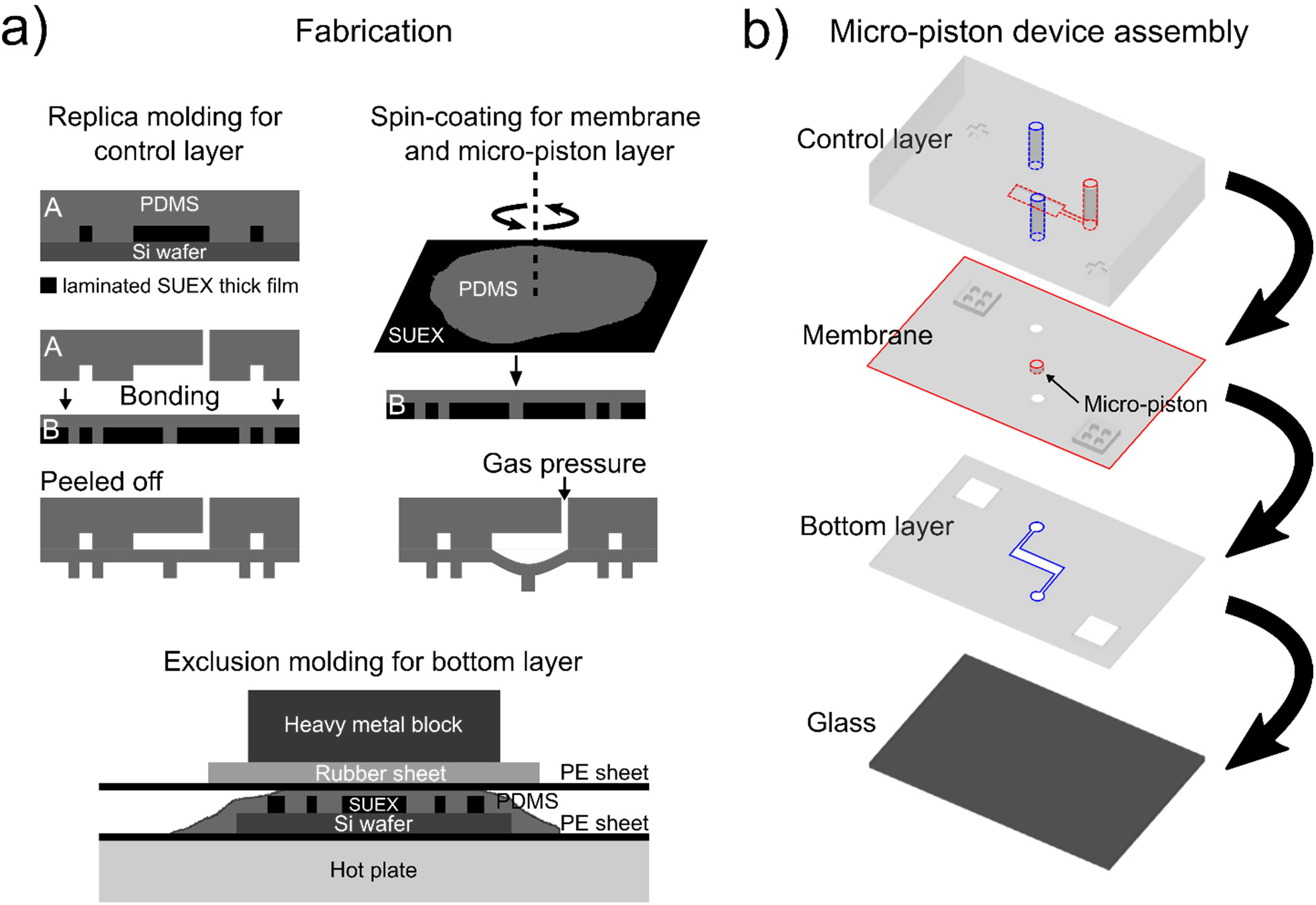
Schematic of the fabrication process and assembly of the micro-piston device. (a) Fabrication of the micro-piston device which is mainly composed of PDMS by different forms of soft lithography: standard replica molding for control layer; spin-coating for membrane and micro-piston layer; and exclusion molding for bottom layer. (b) Micro-piston device is formed by the layers shown in the 3D model by the assembly of control layer, membrane and micro-piston layer, bottom layer open channels and glass substrate, respectively.

**Supplementary Figure S2.**
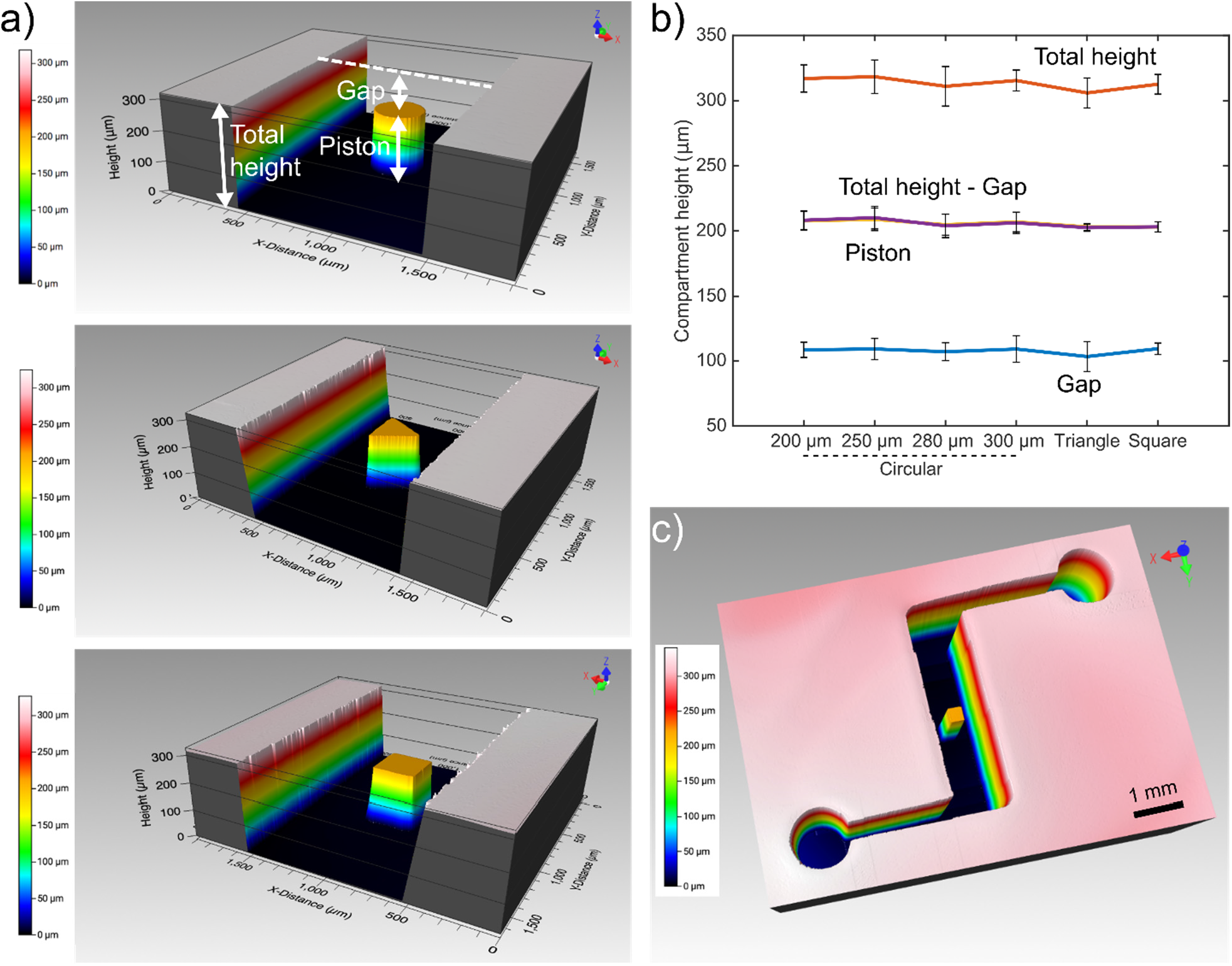
Micro-piston device characterization and average heights of the compartments. (a) 3D optical profilometer image of the micro-piston device showing the compartment including total height of the channel where the micro-piston is suspended, height of the micro-piston and the gap between the piston and surface of the channel. (b) The compartmental heights were measured using optical profilometry for all the devices used in cell-culture and mechanical compression experiments (n = 9, 14, 9, 20, 6 and 6 devices for 200, 250, 280, 300 μm diameter circular, triangle and square micro-piston devices, respectively). For biological cell experiment settings, ‘Gap’ is the distance between the bottom of the micro-piston suspended above the cells and the glass surface on which cells are cultured in static condition of the device. (c) 3D optical profile of one piston per cell-culture chamber is shown as a stitched scan consisting of a series of overlapping individual measurements on a large area.

**Supplementary Figure S3.**
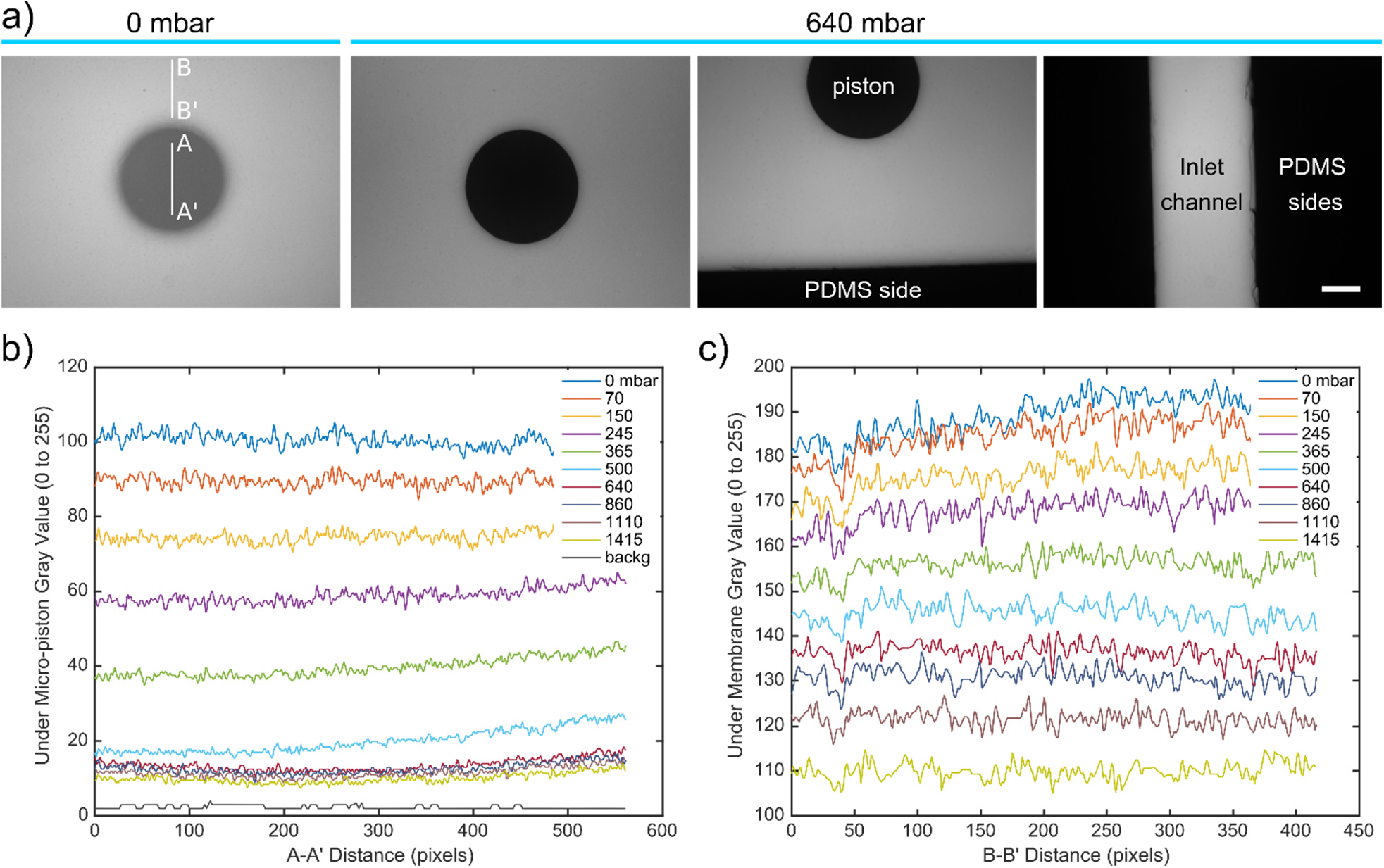
Green fluorescent protein (GFP) solution displacement by membrane deflection in micro-piston device under conditions corresponding to the cell compression experiments. (a) GFP image under the micro-piston while the device was in static stage at 0 mbar and GFP images, including dark regions where there was no GFP solution present, such as under the micro-piston, while the device is in compressed form at 640 mbar. PDMS channel sides and the inlet channel region with GFP solution are also shown (Scale 100 μm). (b) Gray values plotted versus distance, calculated based on the ROI of the GFP solution under the micro-piston compressed at different pressures from static state (0 mbar) to 1415 mbar. The background calculated from the PDMS side channels on the images in (a) is added for reference. (c) Plot of gray values versus distance of a representative ROI of the GFP solution under the membrane deflected at different pressures from static state (0 mbar) to 1415 mbar. Background gray value in (b) is valid for (c) in the same device.

**Supplementary Table T1.** Height versus mean gray value ratios of static to compressed (640 mbar) form of devices, derived from micro-piston device height information of 300 μm diameter circular micro-pistons from Fig. S2 and their GFP solution displacements in ROIs by membrane deflection, as shown in the representative ROI given in Fig. S3(c), and calculated as given in Eqn. 1 and Eqn. 2, respectively.

|  | Device 1 | Device 2 | Device 3 | Average |
| --- | --- | --- | --- | --- |
| **Height ratio** | 1.54 | 1.59 | 1.57 | 1.57 |
| **Mean gray value ratio** | 1.43 | 1.36 | 1.39 | 1.39 |

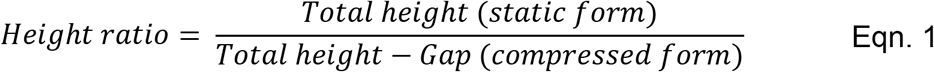

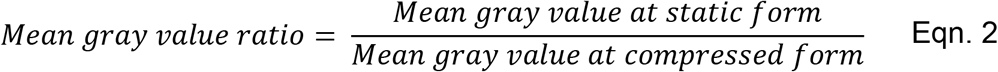

**Supplementary Figure S4.**
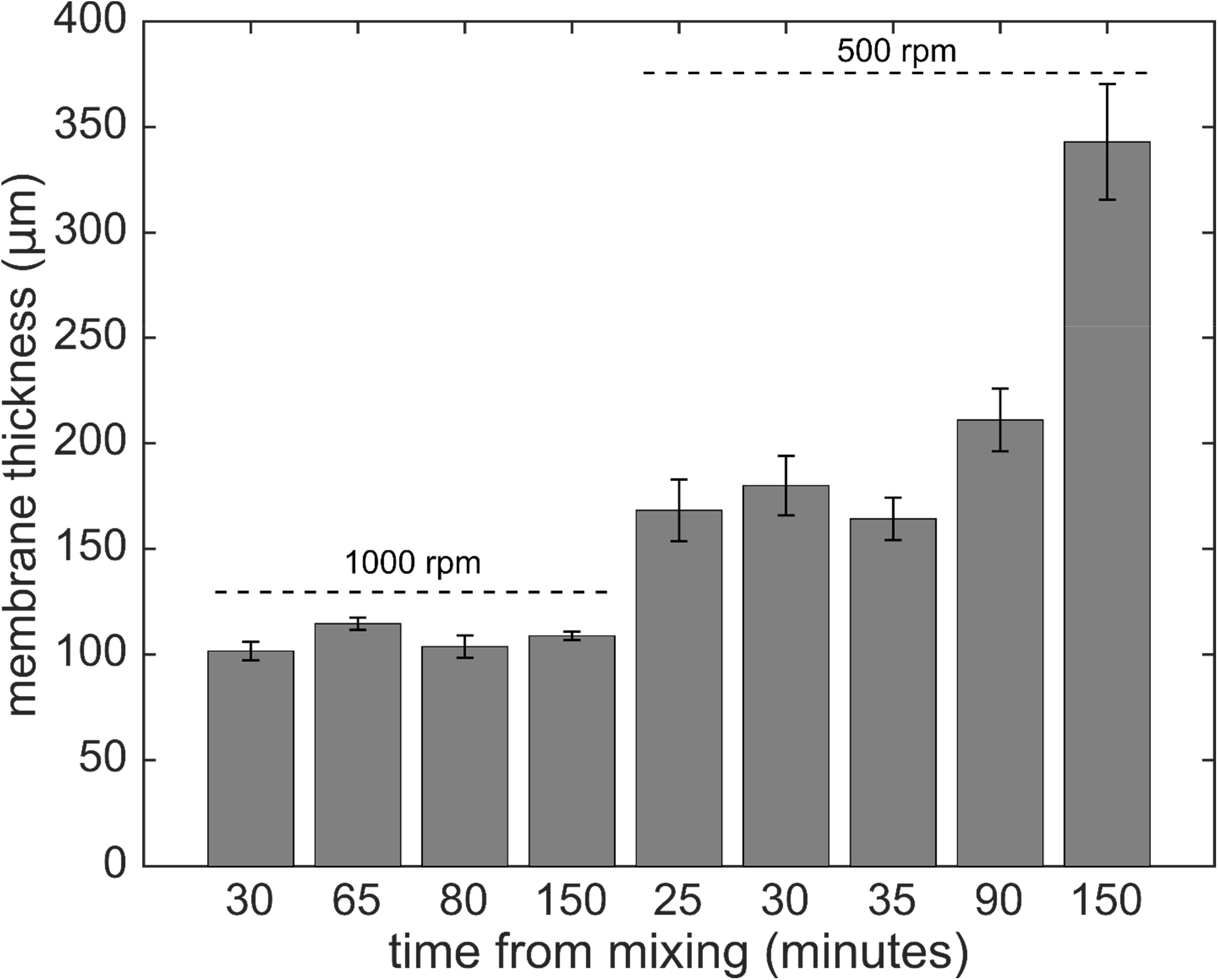
Micro-pistons attached to PDMS membranes of different thicknesses were obtained by varying PDMS spin-coating speeds and vacuum durations from the time of mixing 10:1 PDMS base and curing agent. Spin-speeds were applied for 30 seconds for all membrane/micro-piston samples, either at 500 rpm or 1000 rpm as indicated.

**Supplementary Table T2.** Height measurements for micro-pistons of different shapes on the resist masters and corresponding PDMS molds. Thicknesses of membranes attached to these PDMS micro-pistons before their assembly with other components of the microdevice are also given (as mean ±s.e.m; n = 8, 6 and 6 devices for circular, triangle and square micro-pistons, respectively).

| **Height ($\mu\text{m}$)** | **Circular** | **Triangle** | **Square** |
| --- | --- | --- | --- |
| Micro-piston region in resist master | 214 ±2.4 | 207 ±1.3 | 205 ±1.5 |
| PDMS micro-piston | 214 ±2.6 | 204 ±1.4 | 203 ±1.5 |
| PDMS membrane (thickness) | 210 ±1.7 | 209 ±2.5 | 217 ±2.0 |

**Supplementary Figure S5.**
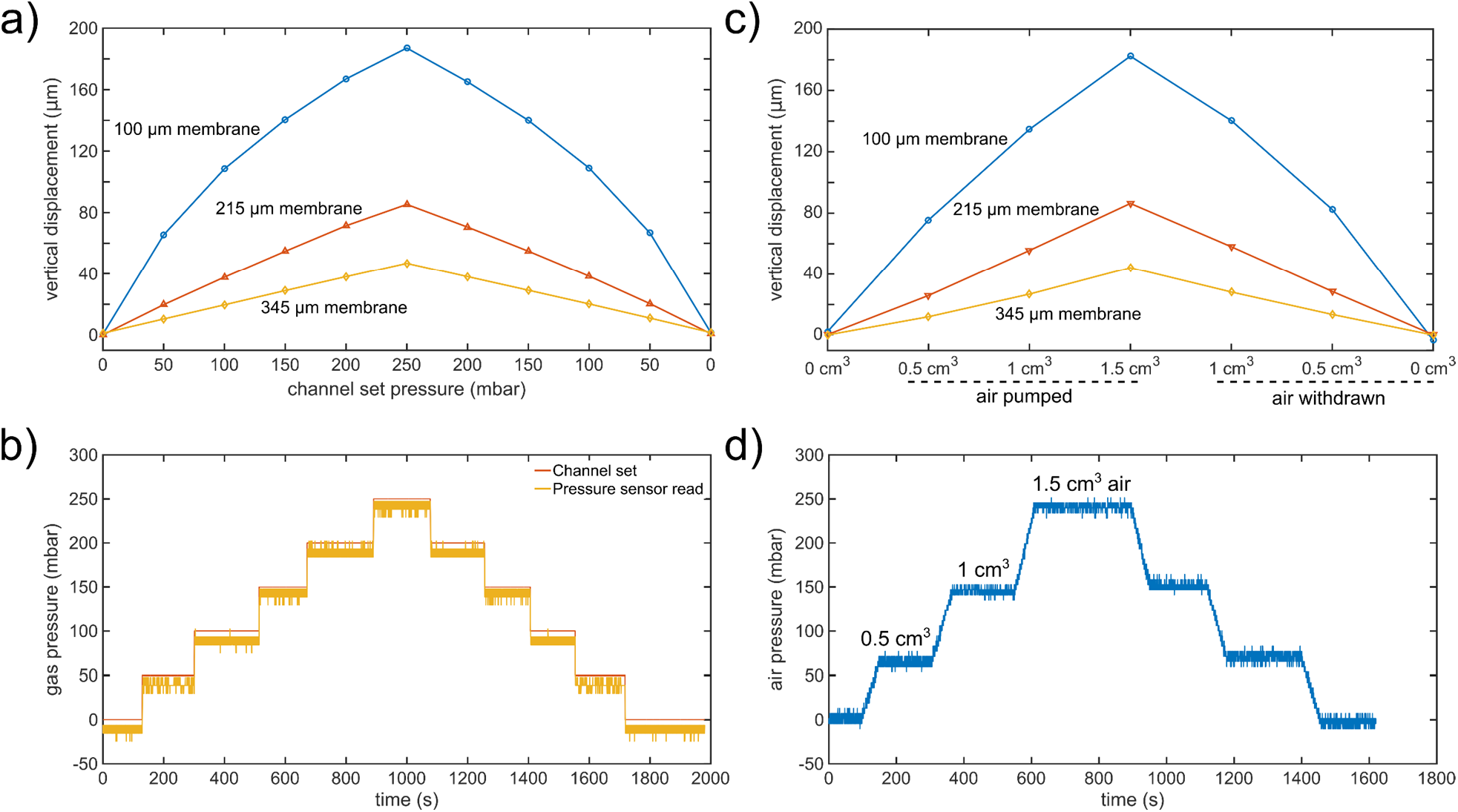
Vertical displacement measurements via optical profilometry to quantify the deflection of 100, 215 and 345 μm thick membranes and micro-piston actuation using two different pressure application systems. (a) Vertical deflection of membranes of different thicknesses as function of applied pressure via a pressure controller system. (b) Corresponding pressure profile for the vertical displacements in (a). ‘Channel set’ is the pressure that was set through the pressure controller of the system while the ‘Sensor read’ is the pressure that was read through the pressure sensor connected within the flow circuit before the gas flow into the control channel of the micropiston device. (c) Vertical deflection of membranes of different thicknesses with various air amounts controlled by a syringe pump system. The corresponding pressure magnitudes are provided in (d).

**Supplementary Figure S6.**
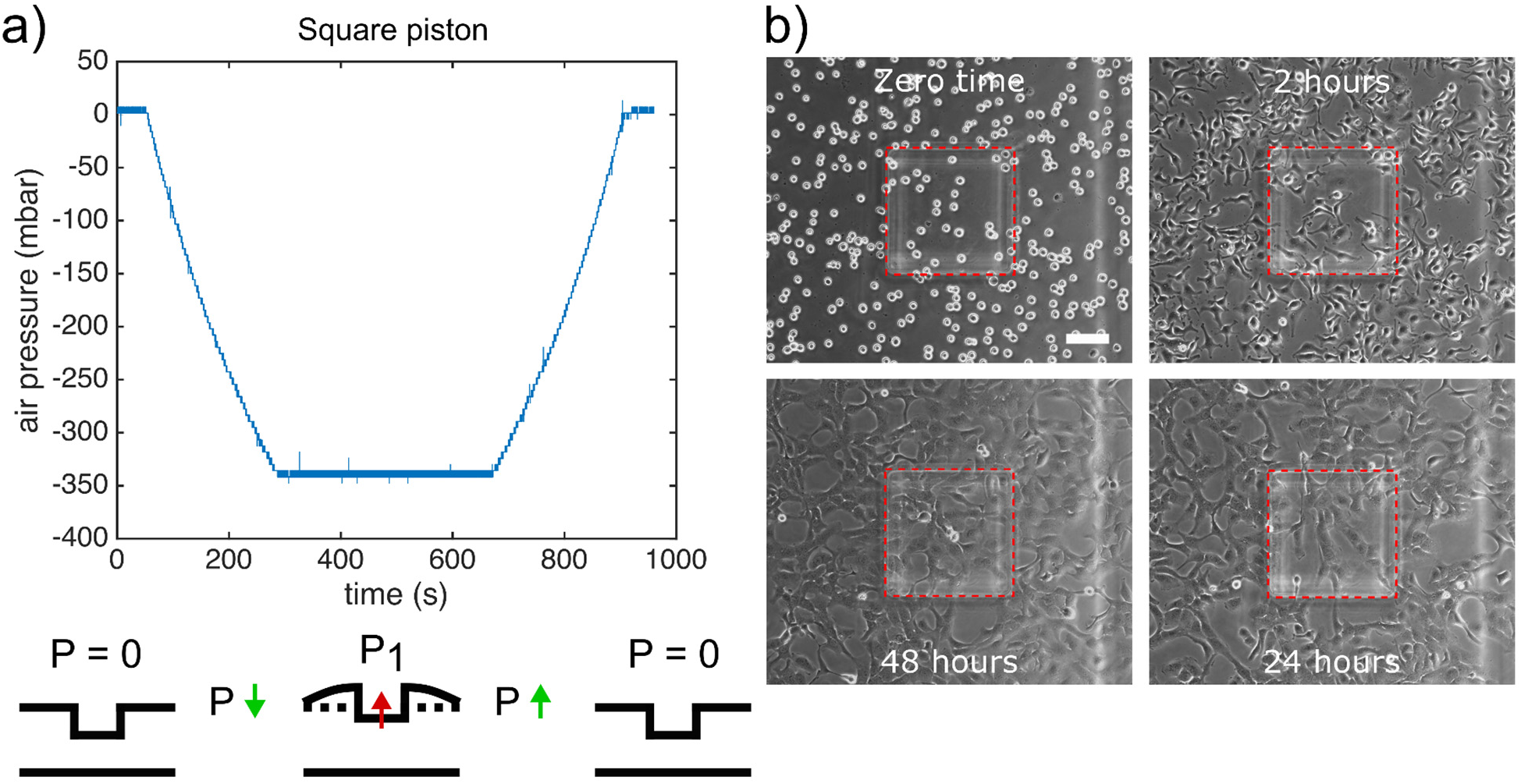
Piston-retracted loading cells into micro-piston device. (a) Pressure sensor read recorded while applying negative pressure to retract the micropiston up into the control microchannel while loading the chamber with cells. After loading and cells settled in, the micro-piston was released to recover its position by applying the same amount of pressure in opposite direction. (b) Representative phase contrast images showing the homogenous cell distribution in piston chamber at zero time of the culture obtained by piston-retracted loading and 2, 24, and 48 hours after the culture growth was started (Scale 100 μm).

**Supplementary Table T3.** Representative data of at least four devices for cell distribution on-chip at zero time of the culture following piston-retracted loading as shown in Fig. S6.

|  | Cell number |  |  | Area (mm<sup>2</sup>) |  |  |
| --- | --- | --- | --- | --- | --- | --- |
|  | Under piston | Around piston | Total in FoV | Under piston | Around piston | Total in FoV |
| **Device 1** | 28 | 232 | 260 | 0.087 | 0.523 | 0.610 |
| **Device 2** | 30 | 263 | 293 | 0.088 | 0.522 | 0.610 |
| **Device 3** | 42 | 307 | 349 | 0.087 | 0.524 | 0.610 |
| **Device 4** | 35 | 248 | 283 | 0.089 | 0.522 | 0.610 |
| **Percentages by average** | **11.4** | **88.6** | **100** | **14.4** | **85.6** | **100** |

**Supplementary Figure S7.**
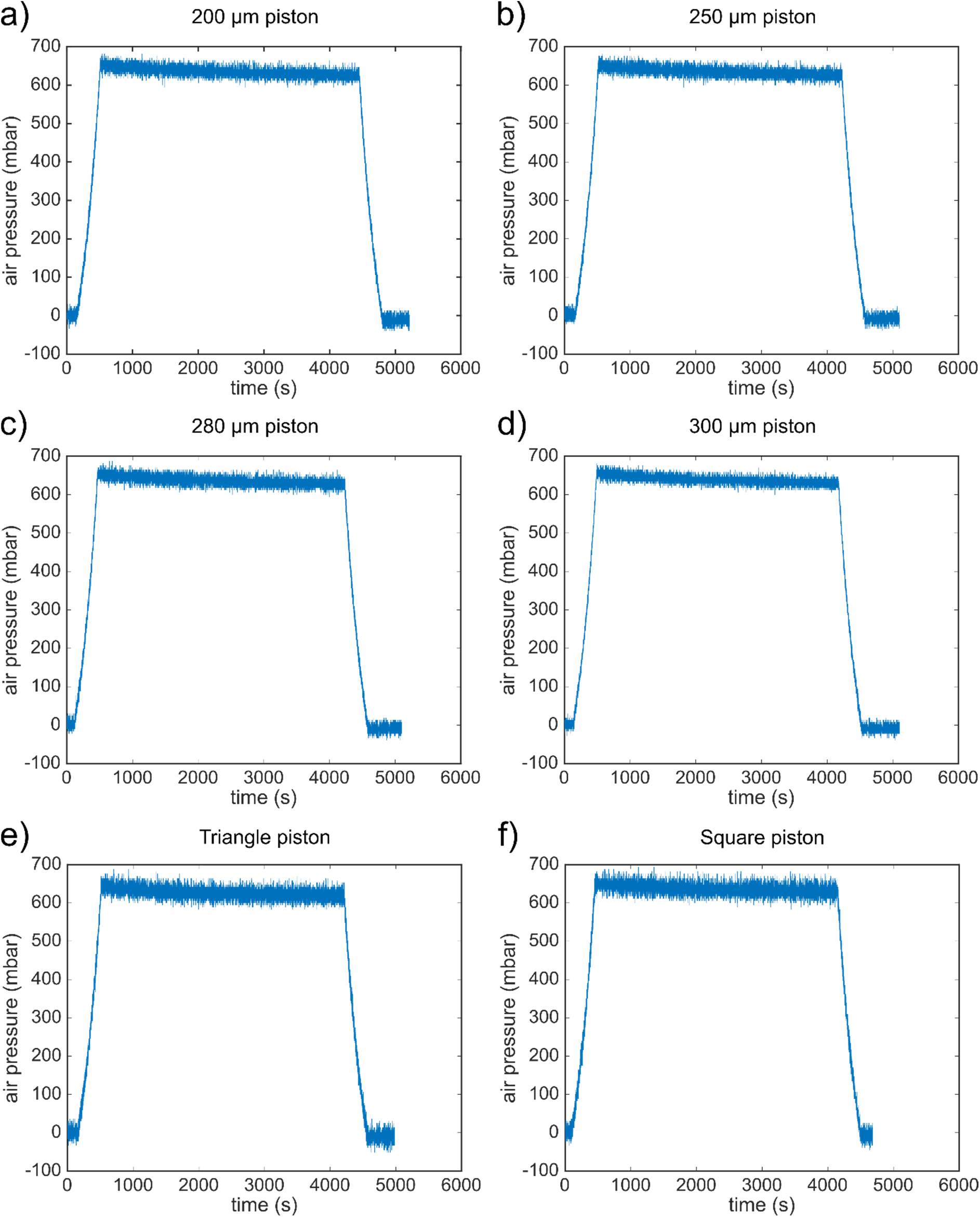
Representative graphs of the pressure sensor readings in the compression experiments on cancer cells with micro-pistons of different diameters for circular shape (a-d) and with different shapes, including triangle (e) and square (f) micro-pistons, run at the same conditions of pressure application (n = 25 devices with dynamic compression).

**Supplementary Figure S8.**
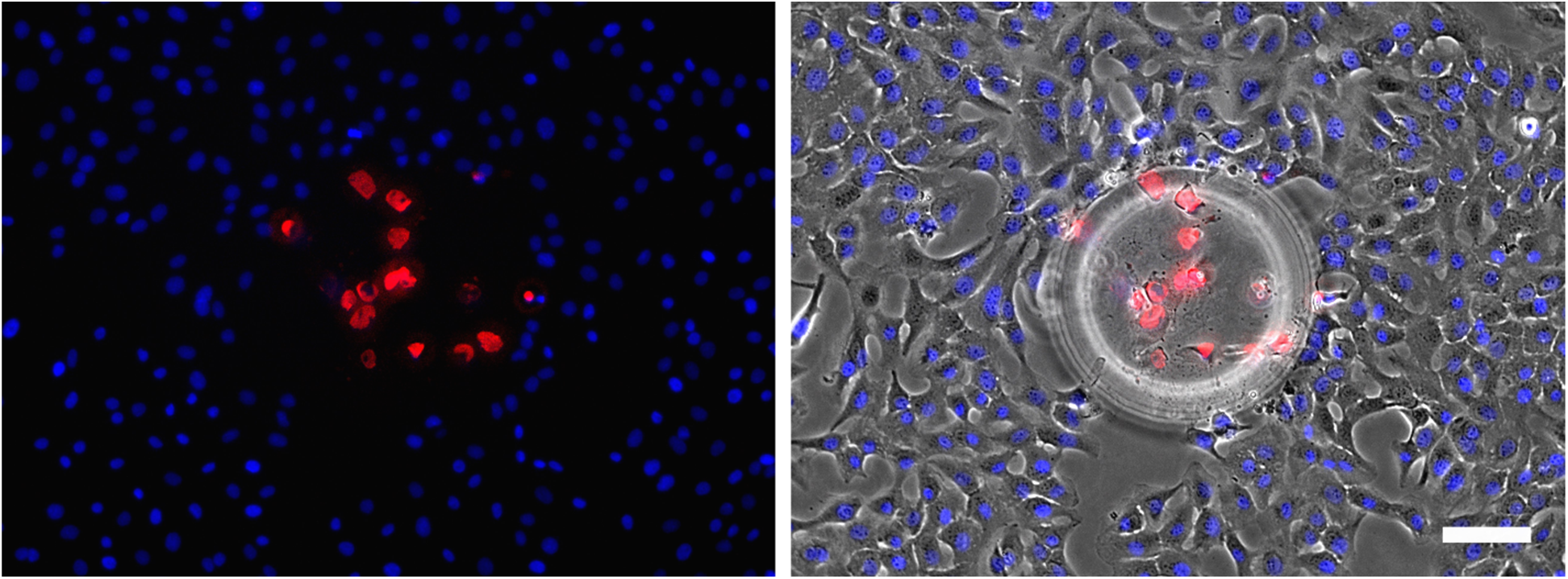
Dead cell and total cell nuclei staining with Ethidium Homodimer (red) and Hoechst dyes (blue), respectively (Scale 100 μm). ImageJ (Fiji) was used for counting cells by analyzing particles on the threshold images. Red: Dead cell nuclei. Blue: Total cell nuclei.

**Supplementary Figure S9.**
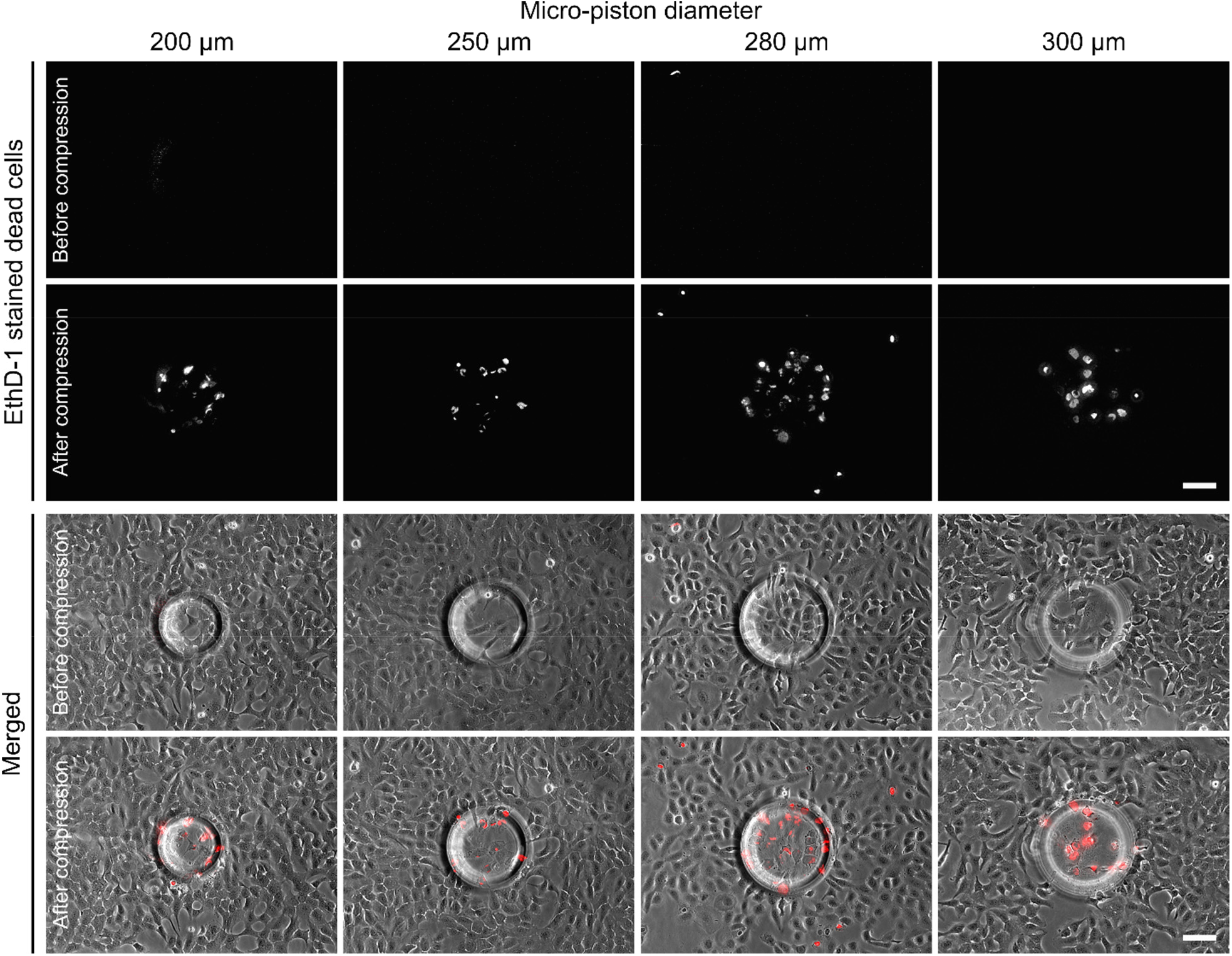
Representative images for cell viability before and after compression applied with various diameter circular micro-pistons (Scale 100 μm). Both control and test experiments were run in the same device. Cells were stained only with 4 μ M EthD-1 (red).

**Supplementary Figure S10.**
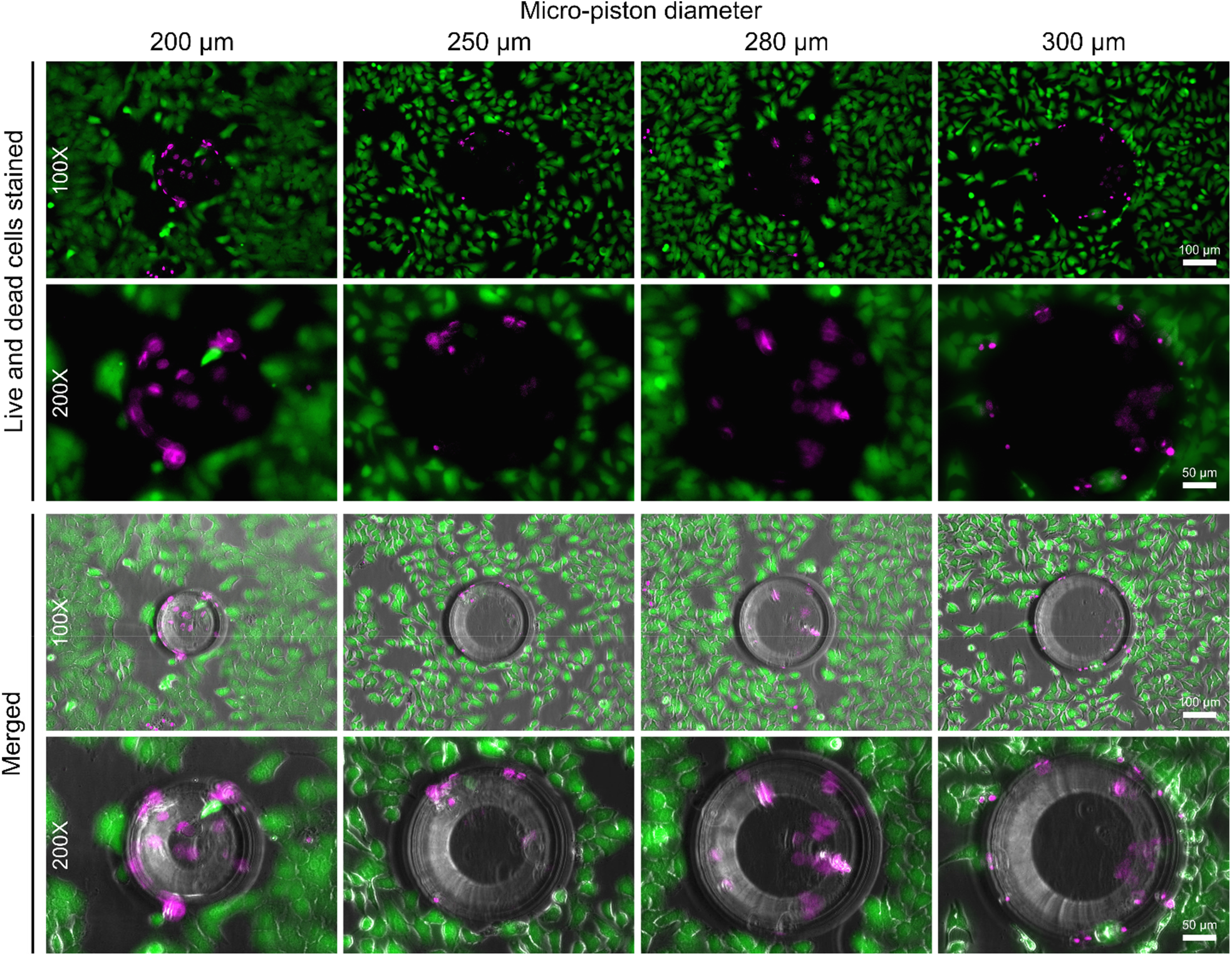
Cell viability after compression applied with various diameter circular micro-pistons. On the row section ‘Live and dead cells stained’, Calcein AM and EthD-1 fluorescence channels were merged. On the row section ‘Merged’, the fluorescence channels were merged with phase contrast. In each, bottom rows are the high magnification images of the micro-pistons shown in the upper rows. Green: Live cells stained with 2 μM Calcein AM, Magenta: Dead cells stained with 4 μM EthD-1.

**Supplementary Figure S11.**
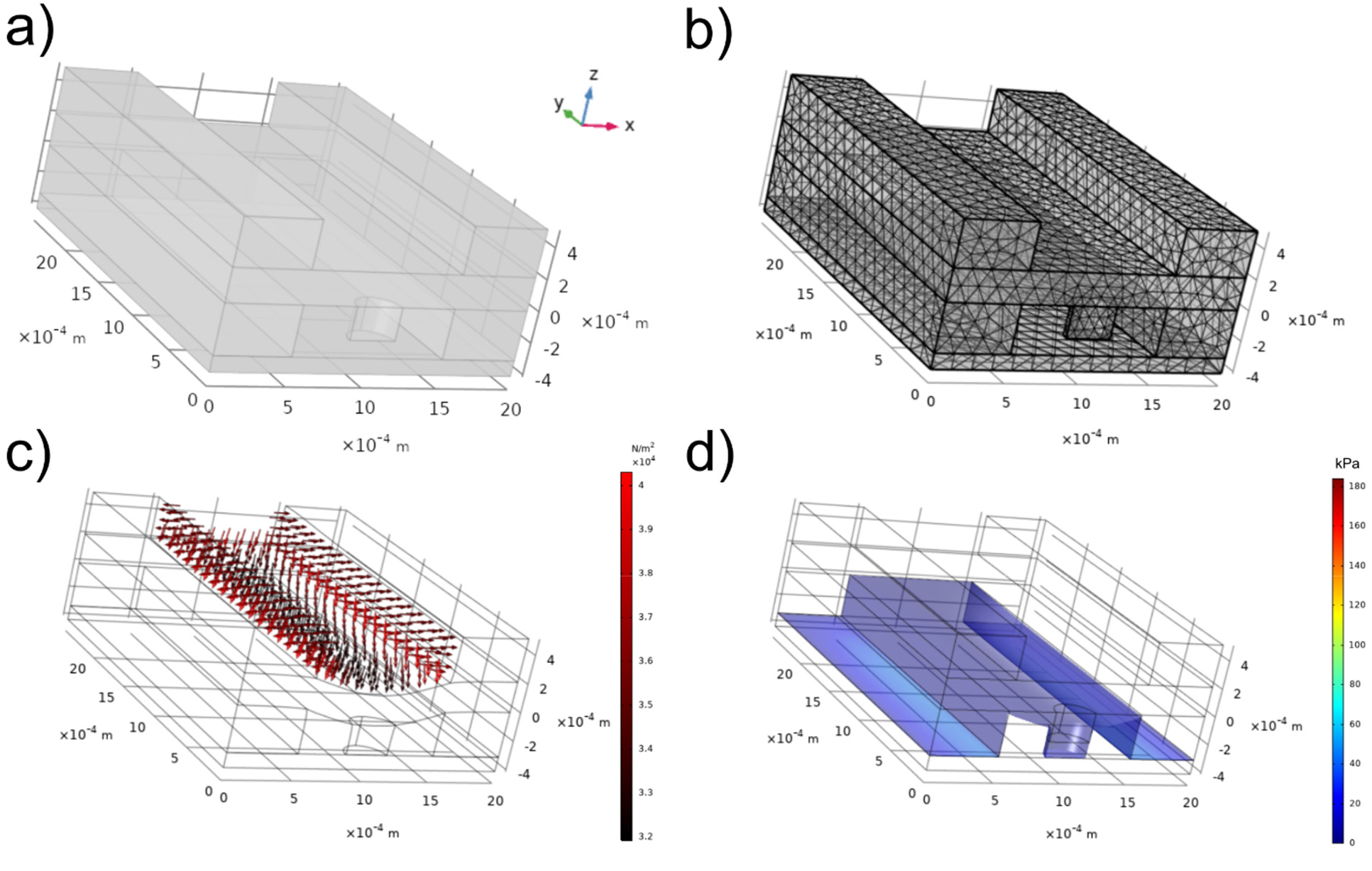
COMSOL simulation results for the actuation of a 300 μm diameter circular micro-piston. PDMS was modelled as hyperelastic material using a Saint Venant-Kirchhoff model. (a) Geometrical model comprising a symmetrical half-cell of the piston and cell-culture chamber. (b) Mesh of the geometry. (c) Example of a 350 mbar boundary load applied to simulate the gas pressure in the upper gas channel. Stationary solutions were obtained with the load varied as parameter ranging from 0 to 700 mbar. (d) Maximum contact pressure at a load of 350 mbar with the deflection of the membrane and micro-piston visualized in relation to the starting position (grid).

### Supplementary Movie Captions

**Supplementary Movie V1**: Green fluorescent protein (GFP) solution displacement frames tracking the micro-piston touch-down for the corresponding applied pressures of 0 mbar to 1415 mbar.

**Supplementary Movie V2**: Bright-field microscopy live-cell image sequence of SKOV-3 cancer cells proliferating inside a PDMS device with 300 μm square micro-piston and orienting under the periphery of the micro-piston over a period of 48 hours. Images were recorded at every 30 minutes in an incubator of 37°C and 5% CO_2_ using a live cell imaging system (CytoSmart). Automated analysis of area coverage (CytoSmart) is shown as inset.

**Supplementary Movie V3**: Compression using the micro-pistons attached to flexible membranes, which were deflected by applying positive pressure from 0 mbar to 640 mbar in 6 minutes. Compression on cells was followed by the retraction of the micro-piston by applying negative (decreasing) pressure from 640 mbar to 0 mbar in 6 minutes. In realtime, each deflection or retraction process for a micro-piston was recorded by time-lapse imaging at 36 frames with 10 sec intervals for the 6-minute period of deflection or retraction.

**Supplementary Movie V4**: Cyclic compression of a SKOV-3 cancer cell monolayer at mild and then higher pressures. In the first pressure profile, a pressure amount ladder was formed up to 354 mbar, during which the cells showed distinct deformation. Subsequently, the cells were chronically compressed for 5 minutes at 354 mbar and rested for 5 minutes at 0 mbar. This cyclic compression was applied for 1 hour. In the following profiles, the pressure was increased to 370, 400, and 640 mbar, respectively, and cells were compressed for up to 2 minutes at each pressure. Time-lapse images were captured once every 10 s from a run of each section of the compression experiment. Cell viability was monitored 10-15 minutes after every pressure application profile was complete, allowing for an incubation time for the viability assay. Live cells (green) and dead cells (magenta) were imaged with Calcein AM and Ethidium homodimer-1 epi-fluorescence, respectively, and merged with phase contrast images.

**Supplementary Movie V5:** Animation of COMSOL Multiphysics simulation results relating to micro-piston actuation and piston contact pressure. Deflection and maximum contact pressure of a 300 μm diameter PDMS piston attached to a 211 μm thick PDMS membrane were simulated and plotted as a function of applied gas pressure. PDMS was modelled as hyperelastic material using a Saint Venant-Kirchhoff model. The Lame parameters λ and μ of PDMS were set at 4.66 MPa and 460 kPa, respectively.

